# Nutritional and host environments determine community ecology and keystone species in a synthetic gut bacterial community

**DOI:** 10.1101/2022.11.24.516551

**Authors:** Anna S. Weiss, Lisa S. Niedermeier, Alexandra von Strempel, Anna G. Burrichter, Diana Ring, Chen Meng, Karin Kleigrewe, Chiara Lincetto, Johannes Hübner, Bärbel Stecher

## Abstract

Microbe-microbe interactions are critical for gut microbiome function. A challenging task to understand health and disease-related microbiome signatures is to move beyond descriptive community-level profiling towards disentangling microbial interaction networks. Here, we aimed to determine members taking on a keystone role in shaping community ecology of a widely used synthetic bacterial community (OMM^12^). Using single-species dropout communities and metabolomic profiling, we identified *Bacteroides caecimuris* I48, *Blautia coccoides* YL58 and *Enterococcus faecalis* KB1 as major drivers of *in vitro* community assembly and elucidated underlying mechanisms of these keystone functions. Importantly, keystone species and bacterial strain relationships were found to strongly vary across different nutritional conditions, depending on the strains’ potential to modify the corresponding environment. Further, gnotobiotic mice transplanted with communities lacking *B. caecimuris* I48 and *B. coccoides* YL58 exhibited morphological anomalies and altered intestinal metabolomic profiles, exposing physiologically relevant functions of these keystone community members. In summary, the presented study experimentally confirms the strong interdependency between bacterial community ecology and the biotic and abiotic environment, underlining the context-dependency and conditionality of bacterial interaction networks.

## Introduction

Symbiotic microbial communities in the mammalian gut are essential for host health. A current challenge in microbiome research is to predict and understand the functional relevance of particular microbial community configurations or “signatures”, associated with health or different diseases (1). Gut microbial community composition is influenced by a variety of abiotic and biotic factors, including temperature, diet, host immune defenses, metabolites and microbe-microbe interactions (2–5). Bacterial interaction networks in particular form the basis of community assembly and structure. However, our understanding of the mechanistic basis underlying these interactions is still incomplete, which greatly limits the functional interpretation of microbiome signatures.

The intestinal microbiota is organized as complex trophic network where individual members engage in cooperative and competitive interactions by nutrient degradation, exchange of metabolites and production of inhibitory compounds (6). Pair-wise interactions are often influenced by the presence of one or more other species in the community and the resulting higher-order interactions limit the predictability of models based on pair-wise interactions alone (7, 8). High-throughput sequencing and metabolomics technologies have generated a wealth of data, which can be exploited by modelling approaches to shed light on the underlying processes shaping the microbiome (9, 10). In particular, inference of microbial interaction and co-occurrence networks are powerful tools for delineating microbial community structures (11, 12). While the computationally identified bacterial associations may result from true ecological relationships, they cannot be distinguished from associations occurring due to environmental selection (13). Hence, the biological interpretation often remains uncertain and requires experimental validation (14, 15). For this purpose, synthetic model communities are indicated, where members are well-characterized, interactions can be experimentally determined and hypotheses can be verified in a systematic way. These experimental model systems also enable the identification of community members with special importance for the ecosystem, i.e., keystones in Paine’s sense (16), by allowing the implementation of systematic presence-absence studies (17, 18). Microbial keystone taxa are highly connected taxa, which have a major influence on microbiome composition and function at a particular space or time (19). These taxa often, but not always, have an over-proportional influence in the community, relative to their abundance (20). Hence, identifying and targeting keystone taxa may open new entry points for microbiome-targeted therapies.

Here, we aimed to identify keystone members of the Oligo-Mouse-Microbiota (OMM^12^), a gut bacterial model community consisting of twelve species (*Enterococcus faecalis* KB1, *Bifidobacterium animalis* YL2, *Acutalibacter muris* KB18, *Muribaculum intestinale* YL27, *Flavonifractor plautii* YL31, *Enterocloster clostridioformis* YL32, *Akkermansia muciniphila* YL44, *Turicimonas muris* YL45, *Clostridium inocuum* I46, *Bacteroides caecimuris* I48, *Limosilactobacillus reuteri* I49 and *Blautia coccoides* YL58), representing the five major eubacterial phyla of the murine gut microbiota (Fig. 1A) (21). This synthetic community is publicly available, adaptable and stable in gnotobiotic mice (22). In addition, it recapitulates important phenotypes of a complex microbiota in mice, including colonization resistance to pathogens (23) and immune development (24). In the past years, the OMM^12^ model has been used by an increasing number of research groups in preclinical disease models, to study gut microbial ecology and evolution and the impact of diet, drugs and phages on the microbiome (25–29). Previous work analyzing mono- and pairwise co-cultures shed light on the metabolic capacity of individual strains and *in vitro* strain-strain interactions in the OMM^12^ community (30, 31). Now, we continued to explore the OMM^12^ interaction network top-down by generating single-species dropout communities aiming to identify keystone species driving community assembly. We analyzed community assembly and strain relationships in different culture media in an *in vitro* batch culture approach, as well as in gnotobiotic mice. In summary, our findings demonstrate a strong environmental context-dependency and conditionality of the keystone species concept and highlight the need for experimental validation of association networks in the relevant biotic and abiotic environment.

**Figure 1:**
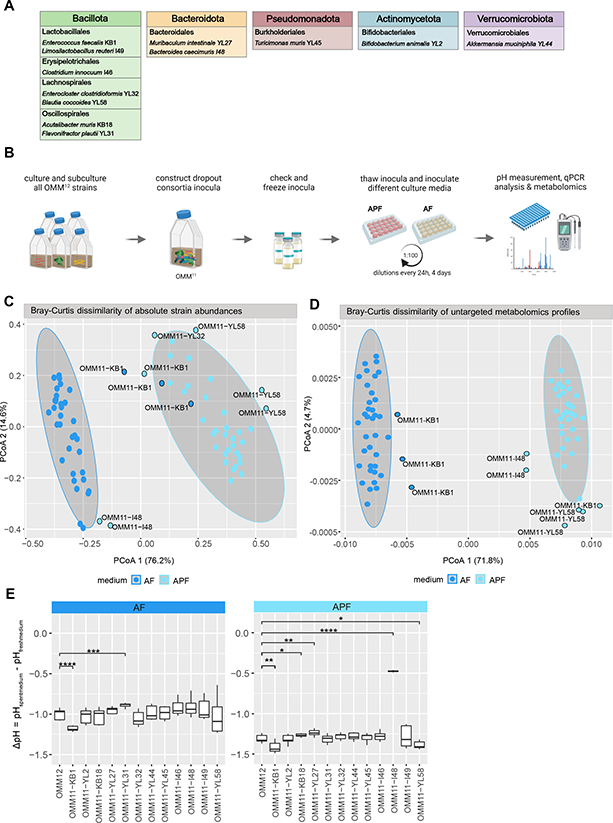
Community assembly of the full consortium and single-species dropout communities in different media. (**A**) Overview of bacterial members of the OMM^12^ consortium. Colored boxes indicate eubacterial phyla represented in the model community: Bacillota (green), Bacteroidota (orange), Pseudomonadota (red), Actinomycetota (blue) and Verrucomicrobia (purple). (**B**) Experimental workflow depicting *in vitro* dropout-community analysis. For the twelve dropout communities and the full consortium bacterial mono-cultures were prepared and inocula mixed accordingly. Generated community inocula were validated and frozen in glycerol stock vials. Thawed inocula were cultured in different media for four days with serial dilutions every 24 h. Culture supernatants from densely grown community cultures (spent media) were sterile-filtered, samples for pH measurements and mass spectrometry were collected and the bacterial pellet stored for qPCR analysis. (**C**) Principle coordinate analysis of community structure in AF and APF medium. Bray-Curtis dissimilarity analysis was performed on absolute abundances of the individual strains (median of three technical replicates) comparing the OMM^12^ and the twelve OMM^11-x^ communities (three biological replicates each) in the two different culture media. Culture media are depicted in different colors. Grey ovals cluster each culture medium with a 95% confidence interval, outliers from the corresponding clusters are highlighted with a black ring. (**D**) Principle coordinate analysis of metabolomic profiles of community SM. Bray-Curtis dissimilarity analysis was performed on untargeted metabolomic profiles of the individual communities comparing the OMM^12^ and the twelve OMM^11-x^ communities (three biological replicates) in the two different culture media AF and APF medium. Culture media are depicted in different colors. Grey ovals cluster each culture medium with a 95% confidence interval, outliers from the corresponding clusters are highlighted with a black ring. (**E**) Environmental modification by community growth in AF and APF. To quantify the environmental modification by the different communities, the pH of the community spent medium (SM) was determined. The ΔpH was calculated by subtracting the SM pH from fresh medium pH. Significant differences between OMM^12^ ΔpH and OMM^11-x^ ΔpH are depicted by asterisks (t-test, p values denoted as *<0.05, **<0.01, ***<0.005, ****<0.001).

## Results

### The carbohydrate environment configures bacterial key species driving community assembly *in vitro*

Previous work using the OMM^12^ *in vitro* community model revealed the strong impact of different media compositions on community assembly (30). Here, to gain insights into the role of the twelve individual strains in community assembly in different nutritional environments, twelve dropout consortia were generated, each lacking one individual strain at a time. Community assembly of all thirteen communities (twelve drop-out consortia, one OMM^12^ full consortium) were studied using a batch culture approach, with slight modifications to the previously published protocol (30). Communities were stabilized for four days with serial dilutions every 24 h (Fig. 1 B) and the absolute abundance of the individual OMM^12^ strains at day four of cultivation was determined by qPCR as normalized 16S rRNA copies per ml culture. The chosen rich culture media differ in the supplied carbohydrate sources: AF medium contains glucose as the major carbon source, while APF medium was supplemented with different mono-(arabinose, lyxose, xylose, rhamnose and fucose) and polysaccharides (mucin, xylan and inulin).

Of note, while the species richness was the same in both media, with eight of twelve bacteria of the consortium coexisting after four days of cultivation, differences in absolute abundance of the strains were observed between AF and APF media (Fig. S1). Bray-Curtis dissimilarity analysis of the median absolute abundance of the individual strains in all communities revealed clear differences (95% confidence interval) between communities grown in AF medium and APF medium (Fig. 1C). Notably, three dropout communities stood out from the corresponding clusters: the community lacking *E. faecalis* KB1 in AF medium and the communities lacking *B. caecimuris* I48 and *B. coccoides* YL58 in APF medium (Fig. 1C). This suggests that depending on the nutritional environment, different bacterial strains take on a key ecological role in driving community assembly.

### Altered community assembly is linked to distinct environmental modifications

We hypothesized that the influence of an individual strain on community assembly is connected to changes in the metabolic environment. To test this, we analyzed spent culture media by untargeted metabolomics. Thereby, a list of unique metabolomics features was curated and compared between fresh and spent media (SM) of the corresponding communities. All samples from communities lacking *E. faecalis* KB1 in AF medium and *B. caecimuris* I48, as well as *B. coccoides* YL58 in APF medium showed distinct metabolomic profiles compared to the other communities in the respective media by both, Bray-Curtis dissimilarity analysis (Fig 1D) and hierarchical clustering of significantly changing metabolites (Fig. S2). This indicated that the strains’ key role in community assembly is strongly connected to their abilities to modify the chemical environment. Specifically, amino acid production and depletion profiles revealed that only communities including *E. faecalis* KB1 depleted serine and arginine in both culture conditions (Fig. S3). On the other hand, communities lacking *B. caecimuris* I48 in APF medium stood out with lower levels of alanine compared to the medium blank, indicating the importance of this strain for alanine accumulation.

Targeted measurements of SCFA levels in the spent culture media further revealed significantly decreased concentrations of acetic acid, propionic acid, isovaleric acid, isobutyric acid and methylbutyric acid in *B. caecimuris* I48 dropout communities cultured in APF medium (Fig. S3). In contrast, communities lacking *E. faecalis* KB1 cultured in AF medium showed significantly increased levels of propionic acid, methylbutyric acid, valeric acid and isovaleric acid and significantly decreased lactic acid concentrations. Of note, butyric acid levels were strongly depleted in communities lacking *F. plautii* YL31 compared to the full consortium. This is in line with information obtained by genome based metabolic models and monoculture *in vitro* data identifying this strain as butyrate producer (30). Of note, butyric acid levels were also significantly decreased in the dropout community lacking *B. coccoides* YL58, in which *F. plautii* YL31 was as well not detectable (Fig. 2).

**Figure 2:**
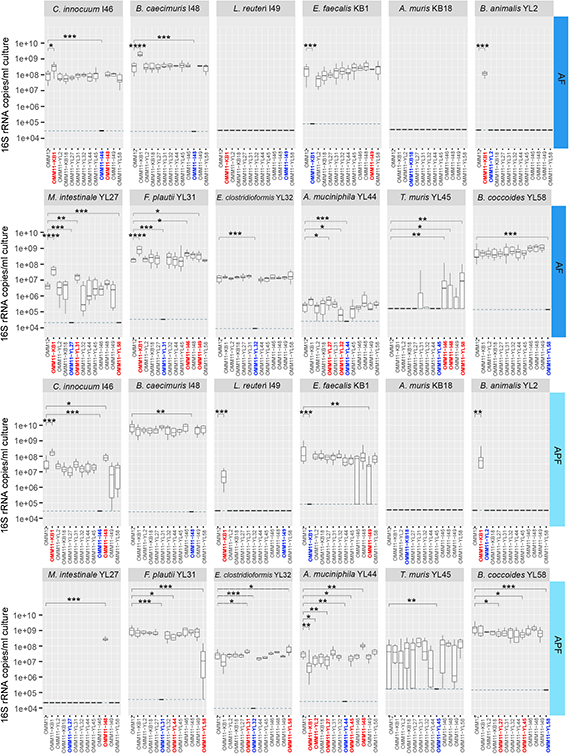
Dissecting strain-strain interactions in the community context using dropout consortia. Community composition of the full consortium and all dropout communities. Absolute abundances of all OMM^12^ strains after four days of dilution in batch culture in AF and APF media were determined by qPCR as normalized 16S rRNA copies per ml culture. Median absolute abundances are shown with the corresponding upper and lower percentile for all individual strains. Significant differences between the full community cultures (N=9) and the dropout communities (N≥6) in the two different culture media are depicted by asterisks (Wilcoxon test, p values denoted as * < 0.05 ** < 0.01, *** < 0.005, **** < 0.0001). Communities in which the absolute abundance of a strain changed significantly are marked in red, corresponding dropout communities are marked in blue. Composition of communities lacking the previously identified keystone species *E. faecalis* KB1, *B. caecimuris* I48 and *B. coccoides* YL58 is additionally shown in Fig. S4 as subset reduced in complexity.

Analysis of pH changes (Δ*pH* = *pH*_*SM*_ − *pH*_*fresh medium*_) in SM revealed that communities lacking *E. faecalis* KB1 showed stronger acidification than the OMM^12^ community in both media conditions (Fig. 1E). In APF medium all communities exhibit more acidic spent medium pH (ΔpH < −1.0) compared to communities in AF medium with the exception of the *B. caecimuris* I48 dropout consortium. Here, a less drastic change in pH was found (ΔpH > − 0.5), indicating that the strong acidification of APF SM is predominantly due to *B. caecimuris* I48.

### Dissecting strain-strain interactions in the community context using dropout consortia

We next compared the absolute abundance of all strains in the corresponding dropout communities to the absolute abundance in the full consortium (Fig. 2). This analysis revealed, that the absence of certain strains have a strong impact on the other strains’ abundances. The strains that influenced the highest number of other strains were *E. faecalis* KB1, affecting five and four strains, *B. caecimuris* I48 affecting one and three strains and *B. coccoides* YL58 affecting two and two strains in AF and APF medium, respectively (Fig. 2, Fig. S4). This is in line with observations made based on Bray-Curtis dissimilarity analysis of community assembly (Fig. 1D).

Specifically, in *E. faecalis* KB1 dropout communities, the abundance of five other species was increased in AF medium. Among those, strains *B. animalis* YL2, *F. plautii* YL31 and *C. innocuum* I46 were previously shown to be inhibited in presence of *E. faecalis* KB1 in co-culture (30). In the absence of *B. caecimuris* I48, three strains were more abundant in APF medium (*M. intestinale* YL27, *A. muciniphila* YL44 and *C. innocuum* I46). Of note, *M. intestinale* YL27 was completely excluded from the consortium in APF medium in the presence of *B. caecimuris* I48 (Fig. 2, Fig. S4). In the absence of *B. coccoides* YL58, two strains were more abundant (*E. clostridioformis* YL32 in AF and *T. muris* YL45 in APF medium), while the abundance of two strains was decreased (*M. intestinale* YL27 in AF and *F. plautii* YL31 in APF medium). Notably, *M. intestinale* YL27 was only detected in the consortium in the presence of *B. coccoides* YL58, indicating a positive dependency of *M. intestinale* YL27 on *B. coccoides* YL58 in the glucose condition but not in APF medium (Fig. 2). Concluding, this underlines that different community members take on a keystone role in community assembly by strongly affecting the abundance of several other strains, depending on the nutritional environment: *E. faecalis* KB1 in AF medium and *B. caecimuris* I48 and *B. coccoides* in APF medium.

### The keystone function of *B. caecimuris* is dependent on the availability of polysaccharides

The pronounced effect *B. caecimuris* I48 had on community assembly in the APF medium, paired with the observation of strongly decreased spent culture medium pH and altered metabolic profiles, suggested a strong interdependency between the availability of polysaccharides and the ability of this strain to degrade polysaccharides. In line with this hypothesis, xylan and inulin concentrations in spent culture media of communities grown in APF medium were significantly reduced in the full community compared to the consortium lacking *B. caecimuris* I48 (Fig. S5). This was further substantiated by the presence of genes encoding the key enzymes of xylan and inulin degradation (32–34) in the *B. caecimuris* I48 genome (Fig S5, Tab. S1). To confirm that the strong influence of *B. caecimuris* I48 on community assembly and spent media pH is indeed mediated by the presence of polysaccharides, we generated a polysaccharide deficient variant of APF medium (APF^mod^). This medium differs from AF medium as the remaining carbohydrate sources are several C5 and C6 sugars, excluding glucose (Methods). Comparing community composition and absolute strain abundances (Fig. S5) revealed that *B. caecimuris* I48 was significantly less abundant in APF^mod^ compared to APF medium. Moreover, none of the initially altered strains showed significant changes in absolute abundances compared to the full consortium in APF^mod^ (Fig. S5). Furthermore, in the absence of xylan and inulin, *B. caecimuris* I48 did not acidify the culture medium as observed for APF medium (Fig. 3A). This is in line with the observation that specific SCFAs, namely acetic acid, lactic acid and propionic acid, that were significantly decreased in communities lacking *B. caecimuris* I48 in APF medium, were unaltered in APF^mod^ medium (Fig. 3B, Fig. S6). Taken together, this suggests that the keystone role of *B. caecimuris* I48 in APF medium is strongly linked to its ability to consume xylan and inulin.

**Figure 3:**
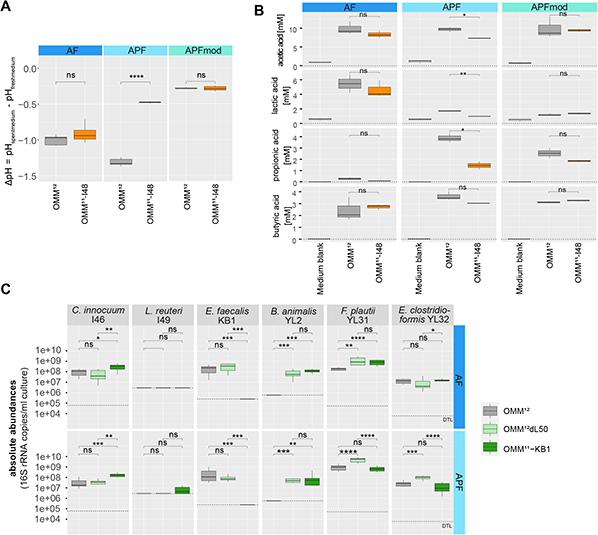
Dissecting mechanisms underlying keystone function of *B. caecimuris* I48 and *E. faecalis* KB1 in the community context. (**A**) Environmental modification by OMM^12^ and OMM^11^-*B. caecimuris* I48 communities grown in AF, APF and APF^mod^ media. To quantify the environmental modification by community growth, the pH of the community spent medium (SM) was determined. The ΔpH was calculated by subtracting SM pH from fresh medium pH. Significant differences between ΔpH of OMM^12^ and OMM^11^-*B. caecimuris* I48 communities are depicted by asterisks (t-test, p values denoted as ns = not significant, **** < 0.001). (**B**) Influence of *B. caecimuris* I48 on SCFA production. Selected SCFA concentrations in community spent media as determined by targeted metabolomics analysis of SM were compared between the full consortium and the OMM^11^-*B. caecimuris* I48 dropout community (t-test, p-value < 0.05 is marked with *, ns = not significant). (**C**) Community composition of the full consortium, the OMM^11^-*E. faecalis* KB1 dropout community and the OMM^12^ community with a *E. faecalis* KB1 ΔL50 mutant strain grown in AF and APF media. Absolute abundances of all OMM^12^ strains after four days of dilution in batch culture were determined by qPCR as normalized 16S rRNA copies per ml culture. Median absolute abundances are shown with the corresponding upper and lower percentile for all individual strains. Significant differences between the full community cultures and the dropout communities (N=9 each) in the different culture media are depicted by asterisks (Wilcoxon test, p values denoted as * < 0.05 ** < 0.01, *** < 0.005, **** < 0.0001).

### *E. faecalis* interactions are multifaceted in the community context

While *E. faecalis* KB1 was previously shown to dominate strain relationships by metabolic interactions in a glucose-rich environment, the strain also inhibited several other community members by bacteriocin production (*B. animalis* YL2, *F. plautii* YL31, *E. clostridioformis* YL32, *C. innocuum* I46 and *L. reuteri* I49) (30). *E. faecalis* KB1 harbors several bacteriocin-encoding loci (30), including enterocin L50, an enterococcal leaderless bacteriocin with broad target range among Gram-positive bacteria (35). We hypothesized that direct interference competition contributes to the capacity of *E. faecalis* KB1 to dominate community assembly. The inhibitory effect of *E. faecalis* KB1 in the community context was most pronounced for *B. animalis* YL2, which is not able to colonize the full OMM^12^ consortium, but can only establish itself in a community lacking *E. faecalis* KB1 in both media conditions (Fig. 2, Fig. S4). A phenotyping approach testing *E. faecalis* KB1 mutants with individual deletions in the three loci for enterocin production (enterocin L50 A and B, enterocin O16 and enterocin E96) identified enterocin L50 A and B as the toxin active against the other community members (Fig. S7).

Next, we performed batch culture experiments to analyze the phenotype of the *E. faecalis* KB1 ΔL50 mutant strain in the community context in AF and APF medium (Fig. 3C, full data set Fig. S8). Interestingly, *B. animalis* YL2 was abundant in the OMM^12^ community including the *E. faecalis* KB1 ΔL50 mutant strain but not in the presence of the wildtype in both nutritional conditions. Further, no significant difference in *B. animalis* YL2 absolute abundance was observed in the *E. faecalis* KB1 ΔL50 mutant community compared to the *E. faecalis* KB1 dropout community, indicating that interaction is mediated by the enterocin. On the other hand, abundance of *C. innocuum* I46 was also decreased in the *E. faecalis* KB1 ΔL50 mutant community, indicating that the observed negative effect of *E. faecalis* KB1 is not due enterocin production, but is mediated by other means such as substrate competition or end product inhibition in both media conditions (Fig. 3C). Interestingly, the effect of the *E. faecalis* KB1 wildtype or the ΔL50 mutant strain on the abundance of *F. plautii* YL31 and *E. clostridioformis* YL32 was not consistent across the two different media, suggesting that the respective changes in absolute abundances of these strains are not solely explainable by interference competition. This highlights that exploitative and interference interactions can occur simultaneously and in a multifaceted fashion.

### Strain interactions are not transferrable across different nutritional environments

As some of the observed strain relationships could not be transferred from one carbohydrate environment to the other, we questioned if testing further nutritional environments may reveal an even higher diversity of bacterial strain relationships in the OMM^12^ consortium. Therefore, the full consortium and three dropout consortia lacking the previously identified keystone species *E. faecalis* KB1, *B. caecimuris* I48 and *B. coccoides* YL58 were cultured in three additional commonly used anaerobic culture media. The chosen media differed in their supplied carbohydrate sources and in the composition and origin of other essential medium components (Tab. S2).

To obtain a quantitative measure for strain relationships (Fig. S9), we defined the measure r_abs_ as the ratio of the absolute abundance of a given strain y in a dropout community lacking strain x vs. the absolute abundance of strain y in the full consortium: 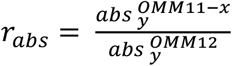. A strain relationship was defined as negative, if the abundance of a strain y was increased in a given dropout community, compared to the corresponding strain abundance in the full community (r_abs_ > 1). A strain relationship was defined as positive, if the abundance of a strain was decreased in a given dropout community, compared to the corresponding strain abundance in the full community (r_abs_ < 1). If a strain y was not detectable in a specific dropout community, but in the full community, the strain relationship was defined as a positive dependency. If a strain y was only detected in a specific dropout community, but never in the presence of the corresponding strain, the interaction outcome was defined as exclusion (Fig. 4A). This analysis revealed a strong variation of strain relationships across different culture media. This even included several cases of contrasting outcomes, as e.g. the relationship between *E. faecalis* KB1 and *A. muciniphila* YL44 (Fig. 4A). Only three strain relationships were found to prevail across nutritional environments: the negative influence of *E. faecalis* KB1 on the strains *B. animalis* YL2 and *C. inoccuum* I46 and the positive relationship between *B. coccoides* YL58 and *M. intestinale* YL27.

**Figure 4:**
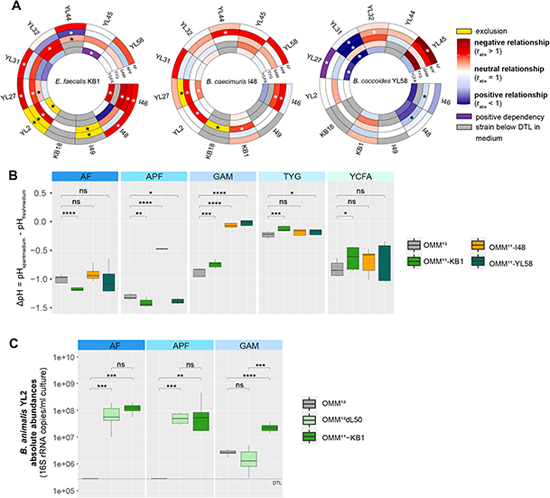
Analysis of strain interactions across different culture media. (**A**) Circular heatmaps depicting strain relationships inferred from changes in absolute strain abundances in the three dropout communities lacking *E. faecalis* KB1, *B. caecimuris* I48 and *B. coccoides* YL58, respectively. Rings depict different culture media, colors indicate the relationship type as determined by r_abs_: negative strain relationship in red, positive strain relationship in blue, exclusion in yellow (strain y only detected in a specific dropout community but never in the presence of the corresponding strain), positive dependency in purple (strain y not detectable in a specific dropout community but in the full community), absolute strain abundance under the detection limit (DTL) in grey. Significantly changed absolute abundances in the corresponding dropout communities compared to the full consortium are depicted by asterisks (Wilcoxon test, p values denoted as * < 0.05). The corresponding raw data set is shown in Fig. S9. (**B**) Environmental modification by community growth in different media. To quantify the environmental modification of the five different media AF, APF, GAM, TYG and YCFA by growth of the full consortium and the three dropout communities OMM^11^-*E. faecalis* KB1, OMM^11^-*B. caecimuris* I48 and OMM^11^-*B. coccoides* YL58, the pH of the community spent medium (SM) was determined. The ΔpH was calculated by subtracting SM pH from fresh media pH. Significant differences between OMM^12^ SM pH and OMM^11-x^ SM pH are depicted by asterisks (t-test, p values denoted as * < 0.05 ** < 0.01, *** < 0.005, **** < 0.0001). (**C**) Absolute abundances of *B. animalis* YL2 in the full consortium, the OMM^11^-*E. faecalis* KB1 dropout community and the OMM^12^ community with a *E. faecalis* KB1 ΔL50 mutant strain grown in AF, APF and GAM media. Absolute abundances of all OMM^12^ strains after four days of dilution in batch culture were determined by qPCR as normalized 16S rRNA copies per ml culture. Median absolute abundances are shown with the corresponding upper and lower percentile for all individual strains. Significant differences between the full community cultures and the dropout communities (N=9 each) in the different culture media are depicted by asterisks (Wilcoxon test, p values denoted as * < 0.05 ** < 0.01, *** < 0.005, **** < 0.0001).

The differential impact of *E. faecalis* KB1, *B. caecimuris* I48 and *B. coccoides* YL58 on community assembly in different nutritional environments was once more reflected in their ability to alter the metabolic environment, as differences in pH modification were observed across the different culture media (Fig. 4B). While e.g. communities lacking *E. faecalis* KB1 show a more acidic spent culture pH in AF and APF medium, less acidification was observed in GAM, TYG and YCFA medium. Interestingly, while the presence of *E. faecalis* KB1 resulted in exclusion of *B. animalis* YL2 in AF and APF medium due to the enterocin production (Fig. 3C), even though still negatively affected, *B. animalis* YL2 coexisted with *E. faecalis* KB1 in GAM (Fig. S9). Again, testing the enterocin mutant strain ΔL50 in the community context in GAM revealed that *B. animalis* YL2 abundance was only significantly increased in the absence of *E. faecalis* KB1 but not in the absence of the toxin only (Fig. 4C). This indicates that in GAM, the enterocin is either not expressed, its effect is overpowered by a simultaneous metabolic interaction or *B. animalis* YL2 is insensitive to the toxin.

### Community assembly and interactions differ across murine gut regions

The observation that community assembly, bacterial interactions and keystone species are strongly dependent on the nutritional environment indicated that OMM^12^ community assembly dynamics and strain relationships might differ across the diverse environments of the murine gastrointestinal tract as well. To test this, germ-free mice (n=8-10 mice per group) were colonized with the full OMM^12^ consortium and three dropout consortia lacking *E. faecalis* KB1, *B. caecimuris* I48 and *B. coccoides* YL58, respectively. After 20 days of colonization, mice were sacrificed and different regions of the gastrointestinal tract (jejunum, ileum, cecum, colon and feces) were sampled for qPCR analysis (Fig. S10, S11). Generally, lower bacterial loads (16S rRNA copies/g content) and higher variability across individual mice were found in the jejunum and ileum compared to cecum, colon and feces (Fig. 5A). Further, especially dropout communities lacking *B. caecimuris* I48 and *B. coccoides* YL58 showed reduced total bacterial loads in cecum, colon and feces compared to mice colonized with the full consortium (Fig. 5A). Bray-Curtis dissimilarity analysis of community profiles, determined by the absolute abundances of the individual strains across different sampling regions, revealed clear differences between community composition in the upper (jejunum and ileum) and in the lower gastrointestinal tract (cecum, colon and feces) (Fig. 5B). While none of the dropout communities stood out from the cluster of the upper gastrointestinal tract (Fig. 5B), the cecal community lacking *B. caecimuris* I48 showed a distinct profile (adjusted p-value 0.0003 Benjamini-Hochberg, Tab. S3), indicating an influential role of this strain in the cecum.

Again, determining the measure r_abs_ to quantify strain relationships from absolute strain abundances over the different gut environments revealed that *B. caecimuris* I48 was mainly positively associated with most other strains in the cecum (Fig. 5C, Fig. S10). This is in line with a particularly strong decrease in total bacterial abundance in the murine cecum in mice colonized with the OMM^11^-*B. caecimuris* I48 community (Fig. 5A). Comparing the relationship of *B. caecimuris* I48 with the other OMM^12^ strains over the different gut regions revealed again contrasting relationships across the different sampling sites (Fig. 5C). In 8 of 10 cases, the strain relationship between *B. caecimuris* I48 and one of the corresponding other strains was opposing. Similar observations were made for the reconstruction of strain relationships for the other two dropout communities, lacking *E. faecalis* KB1 and *B. coccoides* YL58 (contrasting strain relationships in 9 and 6 of 10 comparisons, respectively). Of note, quantifying bacterial strain relationships with *B. coccoides* YL58 revealed a strong positive dependency between *B. coccoides* YL58 and *F. plautii* YL31, as the latter strain seems to be completely dependent on the presence of *B. coccoides* YL58 to establish itself in the murine gastrointestinal tract. A similar relationship, even though not as pronounced, is observed for *C. innocuum* I46, which was as well positively associated with the presence of *B. coccoides* YL58 across all sampling sites.

**Figure 5:**
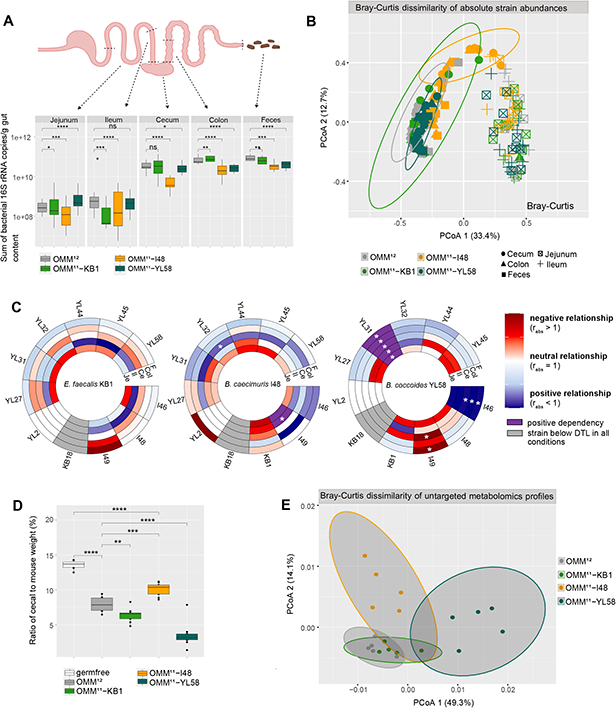
Community assembly and interactions across different regions of the murine gastrointestinal tract. (**A**) Community assembly across different regions of the murine intestine. Germ-free C57Bl/6J mice were inoculated with the full OMM^12^ consortium and the three dropout communities OMM^11^-*E. faecalis* KB1, OMM^11^-*B. caecimuris* I48 and OMM^11^-*B. coccoides* YL58, respectively (N=8-10 mice per group). Absolute abundances of all OMM^12^ strains 20 days after initial inoculation were determined by qPCR as normalized 16S rRNA copies per g gut content and summed up for each mouse. Median sum of absolute abundances are shown with the corresponding upper and lower percentile. Significantly changed sum absolute abundances in mice colonized with the corresponding dropout communities compared to mice colonized with the full consortium are depicted by asterisks (Wilcoxon test, p values denoted as ns = not significant, * < 0.05 ** < 0.01, *** < 0.005, **** < 0.0001). (**B**) Principle coordinate analysis of community structure in different regions on the murine intestine. Bray-Curtis dissimilarity analysis was performed on absolute abundances of individual mice colonized with the OMM^12^ or the three dropout communities OMM^11^-*E. faecalis* KB1, OMM^11^-*B. caecimuris* I48 and OMM^11^-*B. coccoides* YL58 in the different regions of the mouse gut (N=8-10 mice per group). Gut regions are shown in shapes, communities in different colors. Ovals cluster only samples from the murine cecum for each colonization (colors) with a 95% confidence interval. The corresponding statistical analysis is shown in Tab. S3. (**C**) Circular heatmaps depicting strain relationships inferred from changes in absolute strain abundances in the three dropout communities lacking *E. faecalis* KB1, *B. caecimuris* I48 and *B. coccoides* YL58, respectively. Rings depict different sampling regions of the murine intestine, colors indicate the relationship type as determined by r_abs_: negative strain relationship in red, positive strain relationship in blue, positive dependency in purple (strain y not detectable in a specific dropout community but in the full community), absolute strain abundance under the detection limit (DTL) in grey. Significantly changed absolute abundances in the corresponding dropout communities compared to the full consortium are depicted by asterisks (Wilcoxon test, p values denoted as * < 0.05). The corresponding raw data set is shown in Fig. S10. (**D**) Ratio of cecal to mouse body weight of mice colonized with the full consortium or the three dropout communities lacking *E. faecalis* KB1, *B. caecimuris* I48 and *B. coccoides* YL58, respectively. Median cecal to mouse body weight ratios are shown with the corresponding upper and lower percentile (N=8-10 mice per group). Significantly changed ratios in mice colonized with the corresponding dropout communities compared to mice colonized with the full consortium are depicted by asterisks (t-test, p values denoted as ns = not significant, * < 0.05 ** < 0.01, *** < 0.005, **** < 0.0001). (**E**) Principle coordinate analysis of metabolomic profiles of samples from the murine cecum. Bray-Curtis dissimilarity analysis was performed on untargeted metabolomic profiles of cecal samples of mice colonized with the full consortium OMM^12^ or the three dropout communities lacking *E. faecalis* KB1, *B. caecimuris* I48 and *B. coccoides* YL58 (N=5 per group). Communities are depicted in different colors. Grey ovals cluster the corresponding colonization type with a 95% confidence interval.

### Keystone function of *B. caecimuris* and *B. coccoides* is related to altered intestinal physiology and metabolomics profiles in gnotobiotic mice

Mice colonized with communities lacking *B. caecimuris* I48 and *B. coccoides* YL58 exhibited a significantly altered cecal to mouse body weight ratio (Fig. 5D). This ratio is known to be high in germ-free mice and strongly reduced in mice associated with a complex microbiota. For this reason, relative cecal weight is often used as an indicator of the impact of the microbiota on intestinal physiology. Corresponding to the strongly decreased total bacterial loads (Fig. 5A), the relative cecal weight was significantly increased in mice colonized with the OMM^11^-*B. caecimuris* I48 community compared to mice colonized with the full consortium (Fig. 5D). In contrast, even though bacterial loads were decreased in mice colonized with the OMM^11^-*B. coccoides* YL58 community, the relative cecal weight was strongly decreased in those mice compared to the full consortium control group. Of note, this effect was found to be associated with bloating and hardened cecal tissue (Fig. S12).

Next, we analyzed intestinal samples of mice colonized with full and dropout communities using targeted and untargeted metabolomics. In line with the observation that the absence of strains *B. caecimuris* I48 and *B. coccoides* YL58 had a pronounced influence on overall bacterial loads, bacterial community assembly and the host, strongly altered metabolomics profiles were observed for cecal content of mice colonized with communities lacking these species compared to the full consortium control (Fig. 5E). In contrast, cecal metabolomic profiles of mice colonized with the *E. faecalis* KB1 dropout consortium did not differ from the full consortium control. Targeted analysis of SCFAs revealed pronounced differences in SCFA concentrations in the cecal content across mice colonized with the different consortia (Fig. S13). Specifically, a significant decrease in propionic acid, butyric acid, valeric acid, isovaleric acid, isobutyric acid and 2-methylbutyric acid was found in the cecal content of mice colonized with the *B. caecimuris* I48 dropout community compared to mice colonized with the full consortium. Contrasting, cecal content of mice colonized with communities lacking *B. coccoides* YL58 showed significantly increased levels of isovaleric acid and lactic acid, and significantly decreased levels of valeric acid and isobutyric acid compared to the full consortium control. Concluding, we found that *B. caecimuris* I48 and *B. coccoides* YL58 have a particular strong influence on community assembly, total bacterial loads and the metabolomic profile in the cecal content. Most strikingly, the important role of *B. coccoides* YL58 on total bacterial abundance (Fig. 5A) and the host (Fig. 5DE) was not directly inferable from analyzing community composition only (Fig. 5B, Fig. S10, Fig. S11) but became apparent by analyzing physiologic changes in the host and metabolomics profiles.

## Discussion

A central challenge in microbiome research is to move beyond descriptive community-level profiling towards a functional understanding of microbiome signatures. Omics-based approaches have limited traceability of individual community members and experimental means to mechanistically resolve interaction networks. Complementary, the use of synthetic communities provides a reductionist tool to mechanistically resolve interspecies interactions and identify keystone species (17, 18, 36).

Here, we explored the OMM^12^ interaction network using single-strain dropout communities across various culture media conditions and in gnotobiotic mice. This revealed that distinct species drive the community assembly in different environments. Specifically, we identified three environment-dependent keystone species, *E. faecalis* KB1, *B. caecimuris* I48 and *B. coccoides* YL58. Importantly, the kind and extent of how the three species affected community composition differed across culturing conditions and between sampling sites in the murine gut. While *E. faecalis* KB1, a low-abundant member of the mouse microbiome, strongly influenced community assembly *in vitro* by substrate competition and enterocin production, only minor changes in the bacterial abundance of the other community members, metabolomics profiles and physiology of the host were observed in intestinal regions of gnotobiotic mice stably colonized with a *E. faecalis* KB1 dropout community. This difference could be due to the overall low relative absolute abundance of *E. faecalis* in the gut of OMM^12^ mice (30). Increased abundance of *E. faecalis* is found in neonates (37), in antibiotic-treated individuals, graft-versus-host disease or inflammatory bowel disease mouse models, suggesting other environments, where *E. faecalis* potentially takes on a keystone role (38). Our in vitro data suggest, that enterocin-mediated competition is dependent on the nutritional environment, as it was observed in all media conditions except for GAM – and apparently also not in the gut of healthy gnotobiotic mice. Of note, colicin or microcin-dependent competition of *E. coli* and *Salmonella* is also not seen in healthy mice but takes place in the inflamed gut where iron is limiting and both species are highly abundant (39, 40).

In contrast to *E. faecalis* KB1, communities lacking *B. caecimuris* I48 showed distinctly altered metabolomics profiles across culturing conditions and in the murine cecum. *In vivo*, the presence of *B. caecimuris* I48 was positively associated with most other species, whereas *in vitro*, more negative strain relationships with other community members were detected. *B. caecimuris* I48′s keystone role *in vitro* was linked to the presence of the polysaccharides xylan and inulin. Bacteroidetes generally break down dietary polysaccharides outside of their cytoplasm using surface-associated glycoside hydrolases (41, 42). Inulin as supplemented nutrient source was previously also shown in mice to boost *Bacteroides* species and generate altered SCFA profiles (33). Certain *Bacteroides* species also supply inulin breakdown products to other community members by cross-feeding (6). These results suggest that exclusive nutrients enable specific species to take on a keystone role, a concept that might be generalized, but requires further experimental validation.

Most interestingly, the influential role of *B. coccoides* YL58 *in vivo* became only apparent in hindsight of the strongly reduced cecal/body weight ratio and altered cecal metabolomics profiles. From analyzing community assembly in the murine cecum, *B. coccoides* YL58 would not have been inferred as species strongly affecting community structure. Most strikingly, from overall reduced total bacterial abundance in the lower murine gut an increase in cecal to mouse body weight ratio would have been expected; instead it was significantly reduced. The mechanisms underlying the role of *B. coccoides* YL58 in microbiota-host cross-talk will be subject of future work. Besides, *B. coccoides* YL58 was found to be positively associated with several other strains *in vitro* as well as *in vivo*. We hypothesize that this influential role of *B. coccoides* YL58 is linked to its important function as a hydrogen consumer in the OMM^12^ community (30, 43). We reason that increased levels of hydrogen in the absence of this strain could alter the energy balance of hydrogen producing fermentation reactions. In line with this, particularly butyrate producing strains *C. innocuum* I46 and *F. plautii* YL31 are strongly reduced in the absence of *B. coccoides* YL58.

Our previous work showed similarity of OMM^12^ community structure in a polysaccharide supplemented culture medium to that in the murine cecum (30). Nevertheless, we discovered distinct differences in directionality and mechanisms in the underlying strain relationships between the two conditions. Reasons for the observed differences could be manifold, but most likely include the absence of host derived factors, such as antimicrobial peptides (44), oxygen concentration gradients (45) and dynamic pH regulation (46) in the batch culture setup, as well as the lack of structural and spatial heterogeneity, that is present within the lumen of the gastrointestinal tract (47, 48). Hence, while communities similar in compositions can be constructed across different environments, the underlying bacterial interaction networks and therefore resulting overall community functions might differ distinctly.

While we set out to determine universal keystone species of a model gut bacterial community, we found that the keystone species concept is difficult to apply due to its context-dependency and conditionality (20, 49, 50). Overall, we conclude that true “keystone-ness” of a bacterial species that applies across different *in vitro* and *in vivo* conditions is rarely observed. Our data provides an experimental proof that keystone functions of focal microbes can differ in different nutritional and host-associated environments. The presented insights into the sensitivity and dependency of bacterial interactions on the corresponding nutritional environment provided by this and other studies urge the need for a concrete specification of the keystone species concept: a) what key role does a specific strain take on (e.g. shaping the abundance of specific other strains or a concrete metabolic function) and b) in which specific nutritional or chemical environment was this observed (e.g. detailed description of cultivation conditions or sampling region, ideally backed up with additional datasets on physiological and chemical markers).

We conclude that alterations in a community’s interaction network may be overlooked by studying community composition and community-derived correlation and interaction networks. Hence, making use of controllable community models, traceable nutritional environments and a combination of metagenomics and metabolomics approaches is needed to pave the way to elucidate the role of individual species in community functions and delineate general principles of how bacterial interactions shape microbiome function.

## Methods

### Generation of bacterial dropout communities

Bacterial monocultures and subcultures were grown for 24 h each in 10 ml AF medium (30). Dropout community inocula were prepared by first diluting the monocultures in fresh AF to OD600nm 0.1 (BioTek, Epoch2 Microplatereader) under anaerobic conditions. Strains that had an OD600nm < 0.1 were used undiluted. For the generation of each dropout community inoculum, 500 μl of the monoculture dilution or the undiluted monoculture were mixed in a glass culture bottle. Accordingly, eleven different monocultures were used for each dropout community inoculum. The culture bottles with the dropout community inocula were hermetically sealed, discharged from the tent, filled into vials sealed with butyl rubber stoppers (10% glycerol) and frozen at −80°C. Biological replicates were generated from independently prepared monocultures in separately prepared batches of medium.

The following strains were used in this study: *Enterococcus faecalis* KB1 (DSM32036), *Bifidobacterium animalis* YL2 (DSM26074), *Acutalibacter muris* KB18 (DSM26090), *Muribaculum intestinale* YL27 (DSM28989), *Flavonifractor plautii* YL31 (DSM26117), *Enterocloster clostridioformis* YL32 (DSM26114), *Akkermansia muciniphila* YL44 (DSM26127), *Turicimonas muris* YL45 (DSM26109), *Clostridium innocuum* I46 (DSM26113), *Bacteroides caecimuris* I48 (DSM26085), *Limosilactobacillus reuteri* I49 (DSM32035) and *Blautia coccoides* YL58 (DSM26115).

### Culture conditions for dropout community experiments

Bacterial communities were cultivated in 24 well plates (TPP) under anaerobic conditions, thereby diluting the thawed inoculum 1:10 in 1 ml fresh media. Inocula were spiked with 100 μl fresh *M. intestinale* YL27 monoculture, to increase reliability of growth of this strain in the community. Bacterial communities were grown in five different media (Tab. S2): **AF**(30), **APF**(18.5 g.l^−1^ brain-heart infusion glucose-free, 15 g.l^−1^ trypticase soy broth glucose-free, 5 g.l^−1^ yeast extract, 2.5 g.l^−1^ K_2_HPO_4_, 1 mg.l^−1^ haemin, 2.5 g.l^−1^ sugar mix (1:1 arabinose, fucose, lyxose, rhamnose, xylose), 2 g.l^−1^ inulin, 2 g.l^−1^ xylan, 0.025%.l^−1^ mucin, 1 l dH_2_O, 0.5 mg.l^−1^ menadione, 3% heat-inactivated fetal calf serum, 0.5 g.l^−1^ cysteine-HCl·H2O, 0.4 g Na_2_CO_3_), modified **GAM**(Himedia Labs), **TYG**(10 g.l^−1^ tryptone peptone from casein, 5 g.l^−1^ yeast extract, 2 g.l^−1^ glucose, 0.5 g.l^−1^ L-cysteine HCl, 100 ml.l^−1^ 1M K2PO4 pH 7.2, 40 ml.l^− 1^ TYG salt solution (0.05 g MgSO4-7H_2_O, 1 g NaHCO3, 0.2 g NaCl in 100 ml H_2_O), 1 ml.l^−1^ CaCl_2_ (0.8%), 1 ml.l^−1^ FeSO4 (0.4 mg/ml), 1 ml hematine-histidine (pH 8, 0.2M), 1 ml vitamin K3 (1 mg/ml^−1^), 858 ml H_2_O), **YCFA**(10 g.l^−1^ casitone, 2.5 g.l^−1^ yeast extract, 2 g.l^−1^ glucose, 2 g.l^−1^ starch, 2 g.l^−1^ cellobiose, 4 g.l^−1^ NaHCO3, 1 g.l^−1^ L-cysteine, 0.45 g.l^−1^ K2HPO4, 0.45 g.l^− 1^ KH_2_PO_4_, 0.9 g.l^−1^ NaCl, 0.09 g.l^−1^ MgSO4‧7H_2_O,0.09 g.l^−1^ CaCl_2_, 10 mg.l^−1^ hemin, 1.l^−1^ mg resazurin, 100 μl biotin (1 mg/10ml in H_2_O), 100 μl cobalamin (1 mg/10ml in EtOH), 100 μl p-aminobenzoic acid (3 mg/10ml in H_2_O), 100 μl folic acid (5mg/10ml in DMSO), 100 μl pyridoxamin (15 mg/10ml in H_2_O), 1 l dH_2_O, 100 μl Thiamine (5 mg/10ml Stock in H_2_O), 100 μl Riboflavin (5 mg/10ml Stock in H_2_O)). For the polysaccharide deficient variant APF^mod^ of APF inulin, xylan and mucin were left out. Over the total cultivation time of 96 h, 10 μl of the culture were transferred from one well into a new well with 1 ml of fresh medium every 24h (1:100 dilution). For sampling the 24 well plates were discharged from the tent and samples were rapidly processed. For each community culture the full volume (1 ml) was centrifuged at 12,000 rpm for 1 min, the cell pellet was frozen at −20°C and the supernatant was kept for pH measurement and metabolomics analyses.

### pH measurements

pH measurements of bacterial supernatants were performed using a refillable, glass double junction electrode (Orion™ PerpHecT™ ROSS™, Thermo Scientific).

### Metabolomic profiling of bacterial supernatants and cecal content

For metabolomics, 500 μl supernatant or cecal content was transferred to a Spin-X centrifugation tube and centrifuged at 14,000 rpm for 2 min. The membrane was discarded and the flow-through snap frozen in liquid nitrogen.

The untargeted analysis was performed using a Nexera UHPLC system (Shimadzu) coupled to a Q-TOF mass spectrometer (TripleTOF 6600, AB Sciex). Separation of the spent media was performed using a UPLC BEH Amide 2.1×100, 1.7 μm analytic column (Waters Corp.) with 400 μL/min flow rate. The mobile phase was 5 mM ammonium acetate in water (eluent A) and 5 mM ammonium acetate in acetonitrile/water (95/5, v/v) (eluent B). The gradient profile was 100% B from 0 to 1.5 min, 60% B at 8 min and 20% B at 10 min to 11.5 min and 100% B at 12 to 15 min. A volume of 5μL per sample was injected. The autosampler was cooled to 10°C and the column oven heated to 40°C. Every tenth run a quality control (QC) sample which was pooled from all samples was injected. The spent media samples were measured in a randomized order. The samples have been measured in Information Dependent Acquisition (IDA) mode. MS settings in the positive mode were as follows: Gas 1 55, Gas 2 65, Curtain gas 35, Temperature 500°C, Ion Spray Voltage 5500, declustering potential 80. The mass range of the TOF MS and MS/MS scans were 50 - 2000 *m*/*z* and the collision energy was ramped from 15 - 55 V. MS settings in the negative mode were as follows: Gas 1 55, Gas 2 65, Cur 35, Temperature 500°C, Ion Spray Voltage −4500, declustering potential −80. The mass range of the TOF MS and MS/MS scans were 50 - 2000 *m*/*z* and the collision energy was ramped from −15 - −55 V.

The “msconvert” from ProteoWizard were used to convert raw files to mzXML (de-noised by centroid peaks). The bioconductor/R package xcms was used for data processing and feature identification. More specifically, the matched filter algorithm was used to identify peaks (full width at half maximum set to 7.5 seconds). Then the peaks were grouped into features using the “peak density” method (51). The area under the peaks was integrated to represent the abundance of features. The retention time was adjusted based on the peak groups presented in most of the samples. To annotate possible metabolites to identified features, the exact mass and MS2 fragmentation pattern of the measured features were compared to the records in HMBD (52) and the public MS/MS database in MSDIAL (53), referred to as MS1 and MS2 annotation, respectively. The QC samples were used to control and remove the potential batch effect, t-test was used to compare the features’ intensity from spent media with fresh media. For Bray-Curtis dissimilarity analysis of metabolomics profiles features with more than 80% NA values across the analysed samples were removed. The associated untargeted metabolomics data are available on the MassIVE repository (54) with ID MSV000090704.

### Targeted short chain fatty acid (SCFA) measurement

The 3-NPH method was used for the quantitation of SCFAs (55). Briefly, 40 μL of the SM or cecal content and 15 μL of isotopically labeled standards (ca 50 μM) were mixed with 20 μL 120 mM EDC HCl-6% pyridine-solution and 20 μL of 200 mM 3-NPH HCL solution. After 30 min at 40°C and shaking at 1000 rpm using an Eppendorf Thermomix (Eppendorf, Hamburg, Germany), 900 μL acetonitrile/water (50/50, v/v) was added. After centrifugation at 13000 U/min for 2 min the clear supernatant was used for analysis. A Qtrap 5500 Qtrap mass spectrometer coupled to an Exion-LC (both Sciex) was used for the targeted analysis. The electrospray voltage was set to −4500 V, curtain gas to 35 psi, ion source gas 1 to 55, ion source gas 2 to 65 and the temperature to 500°C. The MRM-parameters were optimized using commercially available standards for the SCFAs. The chromatographic separation was performed on a 100 × 2.1 mm, 100 Å, 1.7 μm, Kinetex C18 column (Phenomenex, Aschaffenburg, Germany) column with 0.1% formic acid (eluent A) and 0.1% formic acid in acetonitrile (eluent B) as elution solvents. An injection volume of 1 μL and a flow rate of 0.4 mL/min was used. The gradient elution started at 23% B which was held for 3 min, afterward the concentration was increased to 30% B at 4 min, with another increase to 40% B at 6.5 min, at 7 min 100% B was used which was held for 1 min, at 8.5 min the column was equilibrated at starting conditions. The column oven was set to 40°C and the autosampler to 15°C. Data acquisition and instrumental control were performed with Analyst 1.7 software (Sciex, Darmstadt, Germany).

### Enzymatic assay to determine polysaccharide concentrations

Inulin was measured enzymatically using the Fructan HK Assay Kit (Megazyme Bray, Ireland) according to instructions. Samples were used directly, starting with protocol point E. Assay volume was reduced 20-fold to allow measurements in 96 well plate format.

Xylan was measured enzymatically as xylose after acid hydrolysis using the D-Xylose Assay Kit (Megazyme Bray, Ireland) according to instructions. Xylan was hydrolysed as described in Sample Preparation Example c) in the kit protocol and further measured using the given microplate assay procedure.

### Construction of exchange vectors for the engineering of *E. faecalis*

Vector pLT06 was used for the deletion of enterocins L50A-L50B, Ent96 and O16 in *E. faecalis* strain KB1 (56). A list of all the primers used is provided in Table S4. DNA fragments of 500-1000 bp upstream and downstream of the gene targeted for deletion (homologous arms 1 and 2) were amplified by PCR, using primers to insert restriction sites. *Bam*HI and *Sal*I restriction sites were added to arms 1, and *Sal*I and *Pst*I sites were added to arms 2. The PCR products of the arms were digested using the appropriate restriction enzymes and ligated with pLT06. The ligated products were transformed into NEB 10-beta Competent *E. coli* for propagation and grown on LB plates containing Chloramphenicol (Cm) at 30°C. Colonies were screened for the presence of the inserts using primers pLT06_FW_b and pLT06_RV_b. Positive clones were grown overnight in liquid LB medium containing Cm at 30°C. The plasmids were purified using the PureYield™ Plasmid Miniprep System (Promega) according to the manufacturer’s protocol. The inserts from each construct were sequenced to ensure that no mutations arose during cloning. The resulting exchange vectors were used to generate the respective deletion mutants.

### Construction of *E. faecalis* KB1 enterocin mutant strains

Deletion mutant strains of *E. faecalis* KB1 were engineered by homologous recombination through double cross-over method, as described previously (56). In brief, deletion exchanged vectors were transformed by electroporation into *E. faecalis* KB1. Transformed bacteria were grown on Brain Heart Infusion (BHI) agar plates containing Cm (20μg/ml) and X-Gal (40μg/ml) at 30°C. Blue colonies were inoculated into 5.0 ml of Tryptic Soy Broth (TSB) containing Cm (20 μg/ml) and grown overnight at 30°C. On the next day, the cultures were serially diluted and plated on BHI containing Cm and X-Gal and incubated overnight at 44°C, to force single-site integration by homologous recombination. Light blue colonies were passaged by duplicating them on a BHI plate containing Cm and X-Gal, and then they were screened for the targeted integration using PCR with primers flanking the site of integration. Positive integration clones were grown overnight in TSB with no selection at 30°C. On the next day, the cultures were serially diluted and plated on BHI containing X-Gal only and incubated overnight at 44°C, to force the second site recombination event. The resulting white colonies were passaged by duplicating them on a BHI plate with no selection, and screened for the deletion of the target genes and loss of the plasmid by PCR. Genomic DNA from colonies containing the deletions was amplified and sequenced to confirm gene deletions.

### DNA extraction and purification of *in vitro* and *in vivo* community samples

gDNA extraction using a phenol-chloroform based protocol was performed as described previously (57). First, three small spatula spoons of 0.1 mm zirconia/silica beads (Roth), 500 μl extraction buffer (200 mM Tris-HCl, 200 mM NaCl, 20 mM EDTA in ddH_2_O, pH 8, autoclaved), 210 μl 20% SDS and 500 μl phenol:chloroform:isoamylalcohol (lower phase) were added to the sample. Bacterial cells were lysed with the TissueLyser LT (Qiagen) set on max. speed (50 s-1) for 4 min. Following, samples were centrifuged at 5 min at full speed (14.000 x g). The supernatant was transferred to new 1.5 ml tubes with 500 μl phenol:chloroform:isoamylalcohol. After mixing by inversion the samples were centrifuged again. The supernatant was transferred to new 2 ml tubes containing 1 ml 96% ethanol (p.a.) and 50 μl sodium acetate 3M and mixed by inversion. The samples were centrifuged for min. 30 min at max. speed (14.000 x g) at 4°C. Subsequently, the supernatant was discarded. The pellet was resuspended in 500 μl ice-cold 70% ethanol, mixed by inversion and centrifuged 15 min at max. speed at 4°C. The supernatant was discarded and the pellet was air dried for 5 min. For dissolving, the pellet was resuspended in 150 μl TE buffer (pH 8.0) and stored at 4°C over night. For gDNA purification the NucleoSpin® gDNA Clean-up kit from Macherey-Nagel was used.

### Quantitative PCR of bacterial 16S rRNA genes

Quantitative PCR was performed as described previously (21). 5 ng gDNA was used as a template for qPCR. Strain-specific 16S rRNA primers and hydrolysis probes were used for amplification. Standard curves were determined using linearized plasmids containing the 16S rRNA gene sequence of the individual strains. The standard specific efficiency was then used for absolute quantification of 16S rRNA gene copy numbers of individual strains.

### Genome screening for polysaccharide utilization loci in *B. caecimuris* I48

The genome of *B. caecimuris* I48 was screened for polysaccharide utilization loci (PUL) specific for inulin and xylan degradation found in literature (32, 33) and a PUL database (http://www.cazy.org/PULDB/index.php?sp_name=Bacteroides+caecimuris+I48&sp_ncbi=). Sequences of key enzymes for inulin and xylan degradation were blasted against the *B. caecimuris* I48 genome (https://www.ncbi.nlm.nih.gov/nuccore/CP065319) and names of key enzymes were checked in genome annotations of *B. caecimuris* I48 via word search (“1,4-beta-xylanase”, “beta-xylosidase”, “inulinase” and similar versions). Genome annotations around hits for key enzymes were screened for typical PUL structures (Table S1).

### Spot assays

Bacterial cultures and subcultures were grown for 24 hours each in 10 ml AF medium at 37°C under anaerobic conditions without shaking. Monocultures were diluted to OD_600nm_ 0.1 in fresh AF medium. To generate a dense bacterial lawn, monoculture inocula were diluted in 3 ml LB soft agar to OD_600nm_ 0.01 and poured on an AF medium agar plate. After drying all respective other bacteria were spotted onto the bacterial lawn in duplicates in a volume of 5 μl with OD_600nm_ 0.1. Plates were incubated at 37°C for 24h under anaerobic conditions.

### Mice

All animal experiments were approved by the local authorities (Regierung von Oberbayern and Lower Saxony). Mice were housed under germfree conditions in flexible film isolators (North Kent Plastic Cages) or in Han-gnotocages (ZOONLAB). The mice were supplied with autoclaved ddH_2_O and Mouse-Breeding complete feed for mice (Ssniff) ad libitum. For all experiments, female and male mice between 6-20 weeks were used and animals were randomly assigned to experimental groups. Mice were not single-housed and kept in groups of 2-6 mice/cage during the experiment. All animals were scored twice daily for their health status.

### Mouse experiments

Germ-free C57Bl/6J mice were colonized with defined bacterial consortia (OMM^12^, OMM^11^-*E. faecalis* KB1, OMM^11^-*B. caecimuris* I48, OMM^11^-*B. coccoides* YL58). Mice were inoculated as described previously (22). Mice were inoculated two times (72h apart) with the bacterial mixtures (frozen glycerol stocks) by gavage (50 μl orally, 100 μl rectally). All mice were sacrificed by cervical dislocation at 20 days after initial colonization. Intestinal content from ileum, cecum, colon and feces were harvested, weighed and frozen at −20°C before DNA extraction. Cecal content was sampled in a 2 ml bead beater tube (FastPrep-Tubes Matrix D, MP Biomedical), weighed, snap-frozen in liquid nitrogen and stored at −80°C previous to metabolomics analyses.

### Statistical analysis

For comparison of absolute abundance levels of OMM^12^ strains between experiments, Wilcoxon test was performed using R Studio (version 1.2.5001). Obtained p-values below 0.05 were considered as statistically significant. The vegdist function of the R library *vegan* version 2.5–4 (https://github.com/vegandevs/vegan) was employed to obtain Bray-Curtis dissimilarities between the samples based on absolute abundances. Permutational multivariate analyses of variance (PERMANOVA) were performed in R using the function Adonis (method “bray” with 9,999 permutations). Obtained p values were adjusted using the Benjamini-Hochberg method.

### Data analysis and Figures

Data was analyzed using R Studio (Version 1.2.5001). Heatmaps were generated using the R *pheatmap* package (https://github.com/raivokolde/pheatmap). Plots were generated using the R *ggplot2* package (https://ggplot2.tidyverse.org) and *ggpubR* package (https://github.com/kassambara/ggpubr). Figures were partly generated using BioRender (https://biorender.com) and Affinity Designer (Version 1.10.4.1198).

## Acknowledgements

The authors thank S. Hussain for technical support and members of the Stecher laboratory for helpful feedback and discussions. This research received funding by the German Research Foundation (DFG, German Research Foundation, Projektnummer 395357507– SFB 1371, Projektnummer 279971426 and 315980449), the European Research Council (ERC) under the European Union’s Horizon 2020 research and innovation program (Grant Agreement 865615), the German Center for Infection Research (DZIF) and the Center for Gastrointestinal Microbiome Research (CEGIMIR).

## Author contributions

A.S.W conceived and designed the experiments. A.S.W., L.S.N., A.v.S., A.G.B. and D.R. performed the experiments. A.S.W., L.S.N, C.M. and K.K. analyzed the data. C.M., K.K., C.L. and J.H. contributed materials, strains or analysis tools. A.S.W. and B.S. coordinated the project. A.S.W. and L.S.N. wrote the original draft and all authors reviewed and edited the draft manuscript.

