## Supplementary Material for "Nutritional and host environments determine community ecology and keystone species in a synthetic gut bacterial community"

### Supplemental Information

#### Supplemental Figures

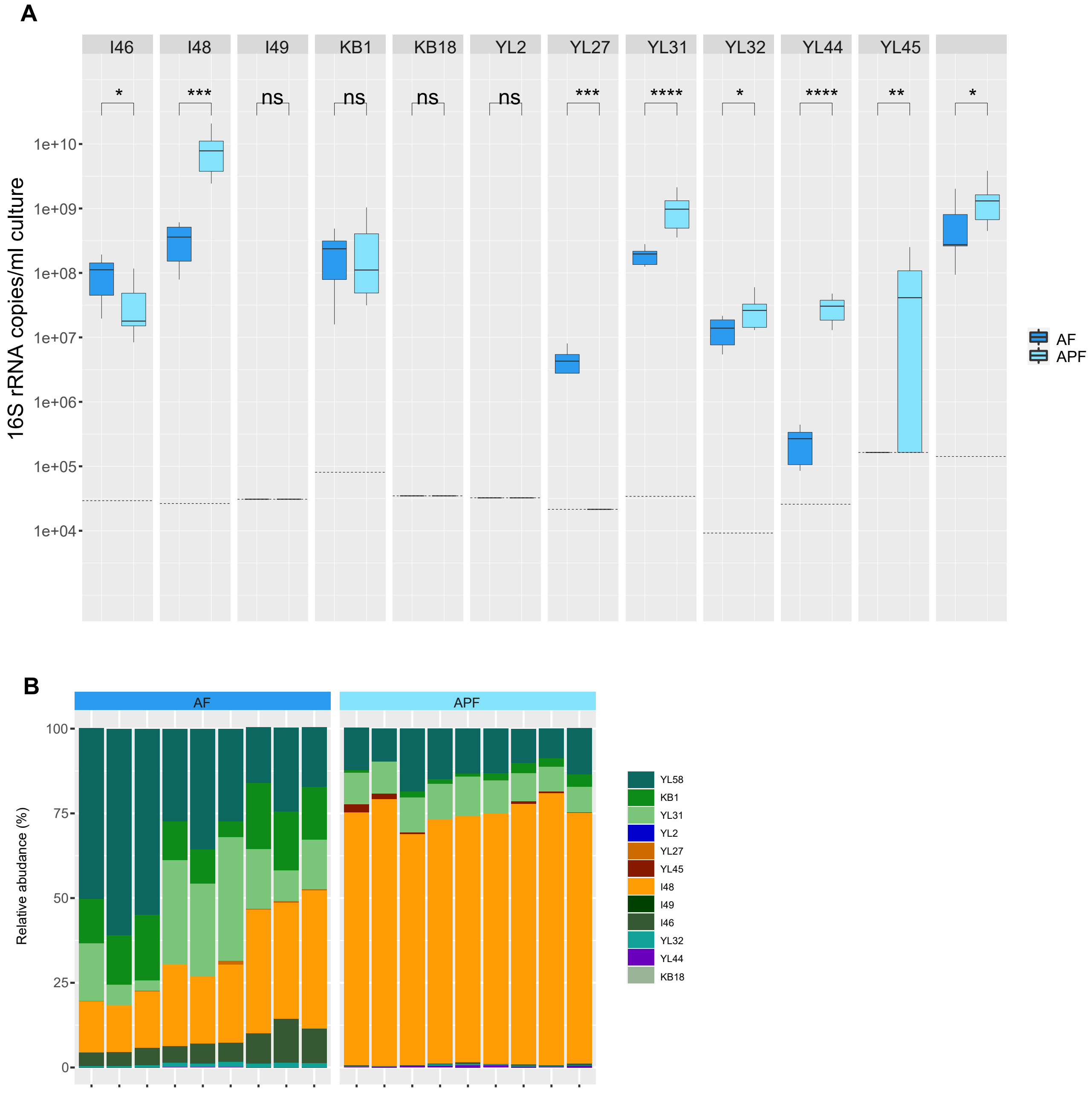


**Fig. S1. OMM$^{12}$ community composition in AF and APF medium.** Absolute abundance of strains (**A**) after four days of serial dilution in batch culture in AF and APF media was determined by qPCR as normalized 16S rRNA copies per ml culture. Median absolute abundances are shown with the corresponding upper and lower percentile for all individual strains. The detection limit for each strain is shown as a dotted line. Strains *L. reuteri* I49, *B. animalis* YL2 and *A. muris* KB18 were below detection limit in the full community context in both culture media. Using a Wilcoxon test, absolute abundances were compared between the communities grown in AF and AFP medium (N=9 each), p values are denoted as * < 0.05, ** <0.01, ***< 0.005, ****< 0.001. Corresponding relative abundances of all replicates were calculated based on absolute abundances (**B**).

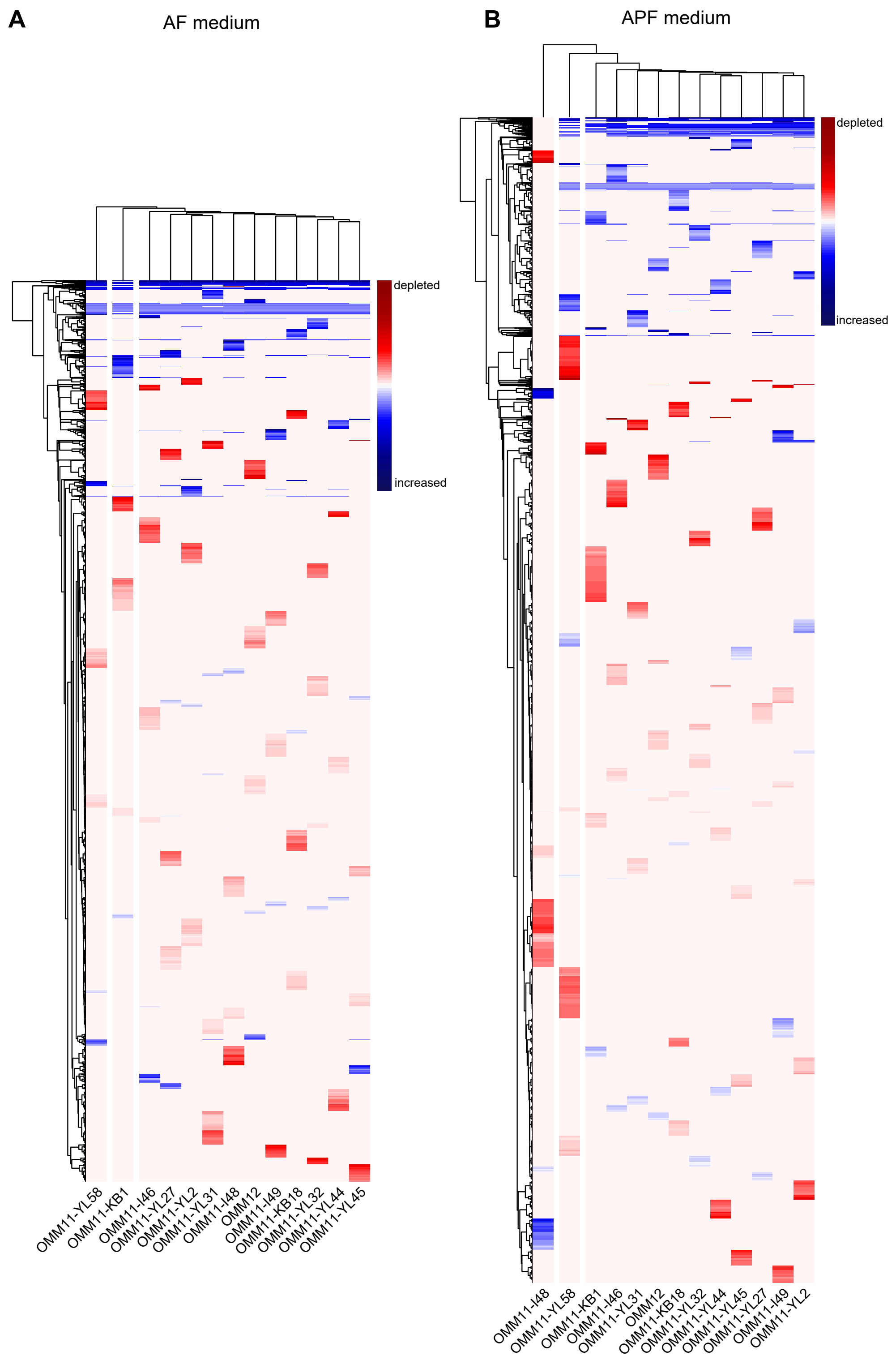


**Fig. S 2. Untargeted metabolomics analysis of community spent media.** Metabolomic profiles after community growth to stationary phase in AF medium (**A**) and APF medium (**B**) were determined by untargeted MS. All metabolomic features that significantly changed in comparison to the medium blank for at least one of the 13 communities are shown (rows, SI data table 2). Levels decreased and levels increased compared to fresh AF or APF medium as determined by the relative foldchange are shown in red and blue, respectively. Hierarchical clustering of community specific profiles revealed more pronounced differences in profiles for communities lacking *B. coccoides* YL58 and *E. faecalis* KB1 in AF medium and the *B. caecimuris* I48 dropout community in APF medium.

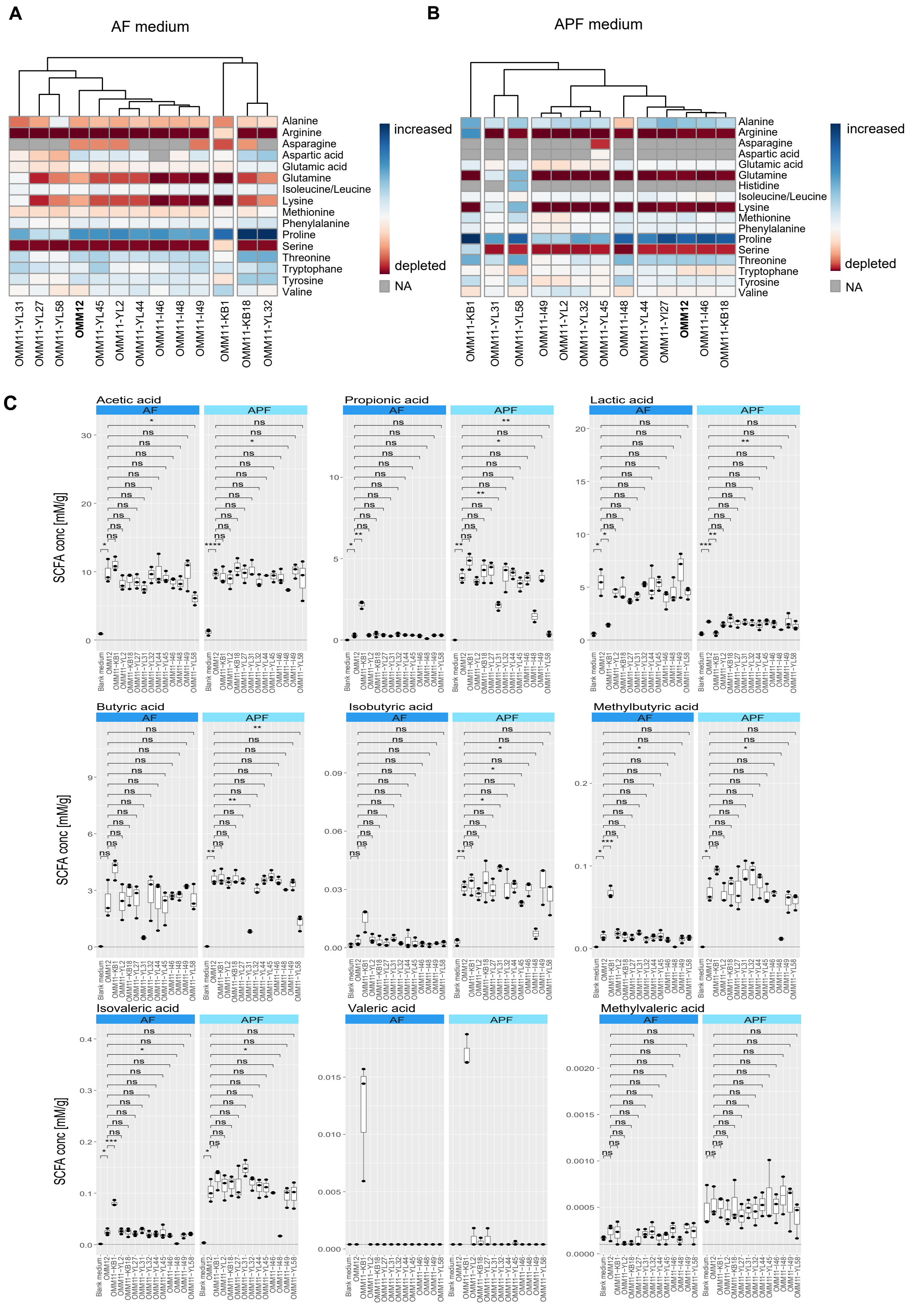


**Fig. S3. Metabolic profiling of community spent media.** Amino acid profiles of community spent media in AF medium (**A**) and APF medium (**B**). Amino acids were annotated by comparing the exact mass and MS2 fragmentation patterns of the measured features to the records in HMBD and the in-house database. Consumption of a specific amino acid from fresh medium is indicated in red, while production in indicated in blue. SCFA concentrations (**C**) were determined by targeted metabolomics analysis of community spent media and fresh media and are shown as median with the corresponding upper and lower percentile. Using a t-test the SCFA concentrations in the dropout communities were compared to the full consortium. p values are denoted as ns = not significant, * < 0.05, ** <0.01, ***< 0.005

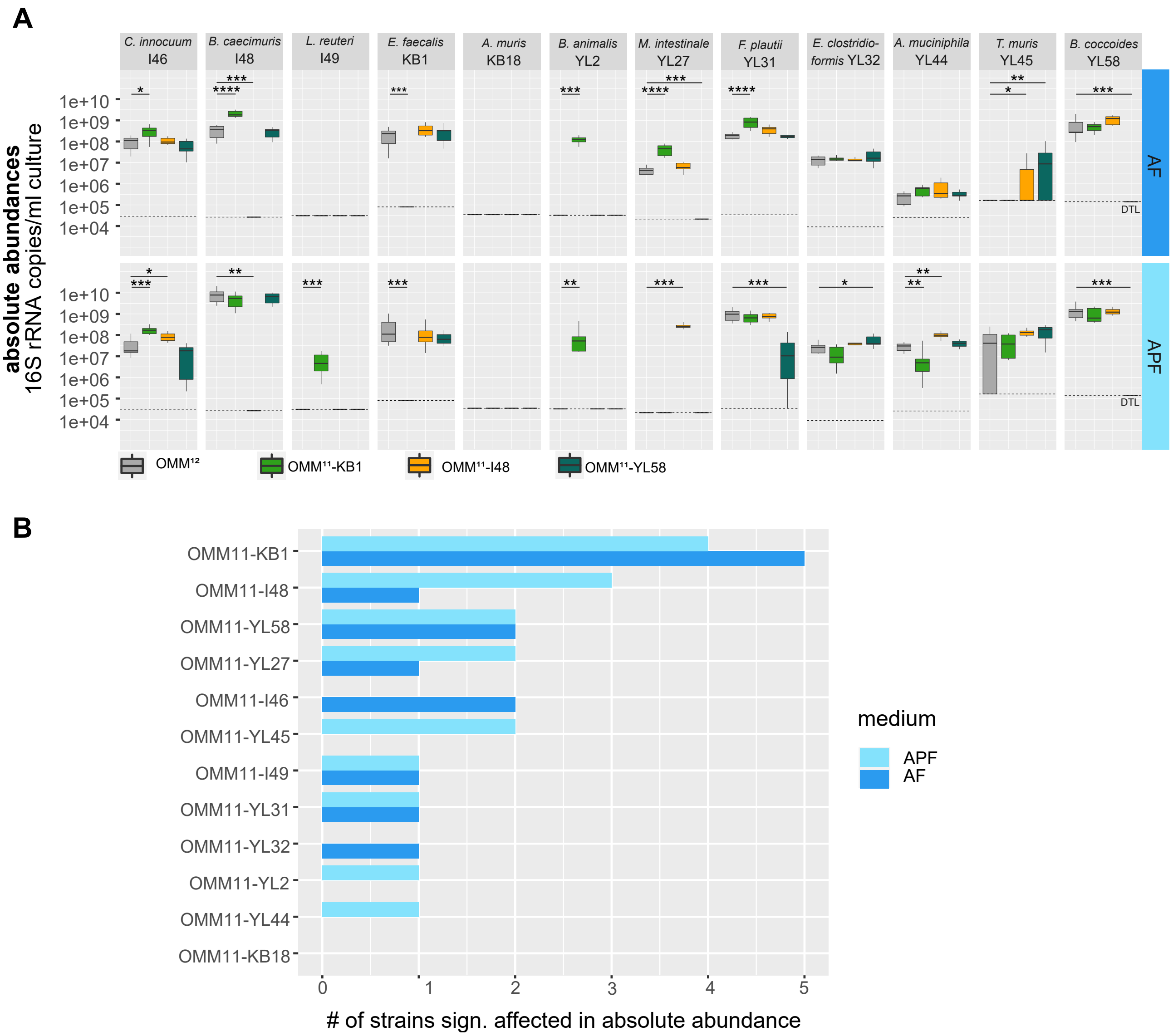


**Fig. S4. Community assembly of OMM$^{12}$ and dropout communities.** Community composition of the full consortium and the three dropout communities lacking *E. faecalis* KB1, *B.caecimuris* I48 and *B. coccoides* YL58, respectively. Absolute abundances of all OMM12 strains after four days of dilution in batch culture in AF and APF media were determined by qPCR as normalized 16S rRNA copies per ml culture. Median absolute abundances are shown with the corresponding upper and lower percentile for all individual strains. Significant differences between the full community cultures (N=9) and the dropout communities (N≥6) in the two different culture media are depicted by asterisks (Wilcoxon test, p values denoted as * < 0.05 ** < 0.01,*** < 0.005, **** < 0.0001) (**A**). To quantify the influence a specific strain had on community assembly, the number of strains significantly affected in their absolute abundance (compared to the full consortium) are shown for the individual dropout communities in a barplot (**B**) for both culture media.

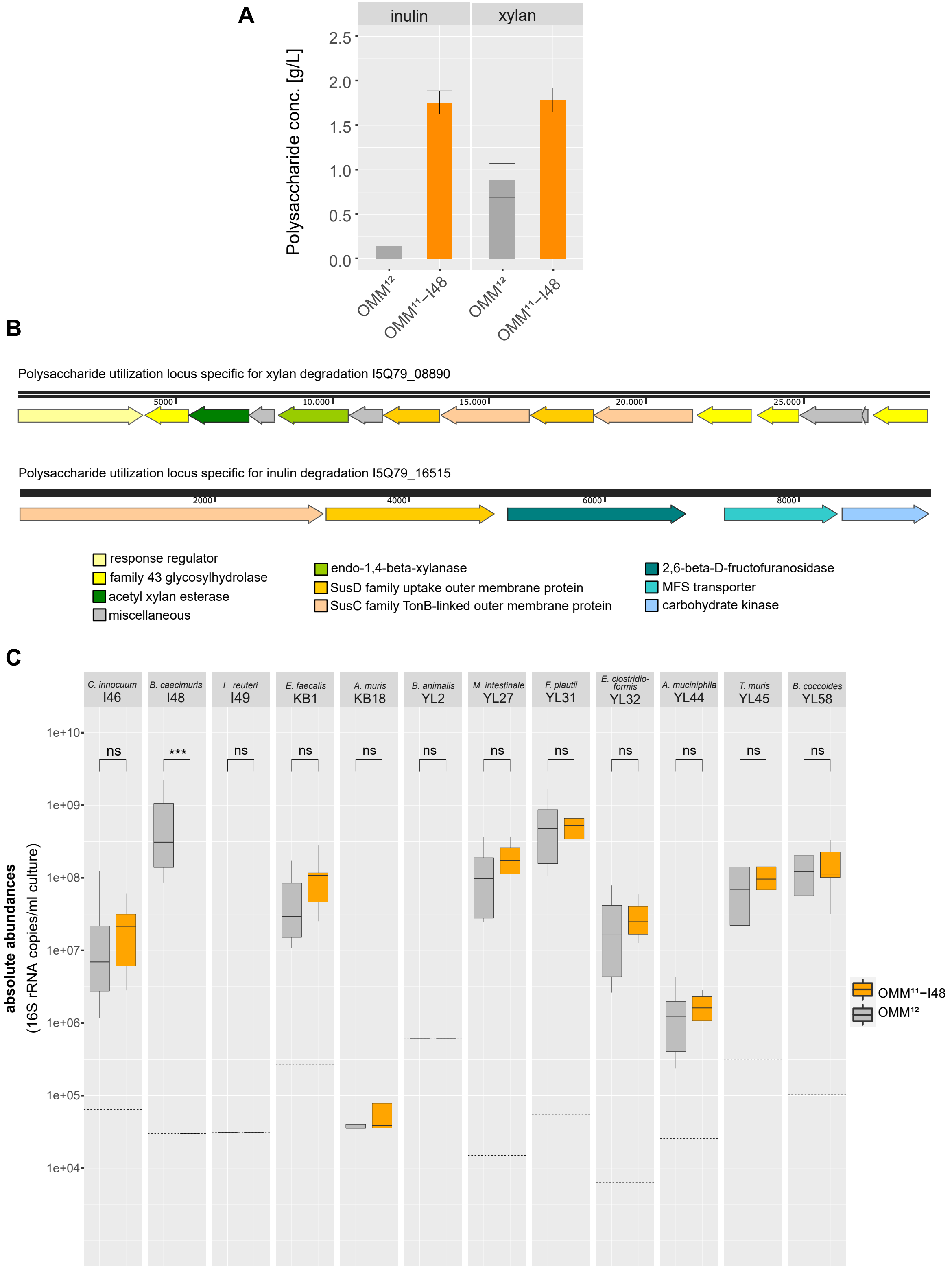


**Fig. S5. Role of polysaccharides for *B. caecimuris* I48 keystone role.** Concentrations of inulin and xylan were measured using an enzymatic assay in community spent medium of the full consortium and the *B. caecimuris* I48 dropout community (**A**). Polysaccharide utilization loci for xylan and inulin as identified in the genome of *B. caecimuris* I48 (**B**). Absolute abundances as normalized 16S rRNA copies per ml culture for the full consortium and a *B. caecimuris* I48 dropout community in APF$^{\mathrm{mod}}$ medium (**C**). Median absolute abundances are shown with the corresponding upper and lower percentile for all individual strains. Using a Wilcoxon test, absolute abundances of the individual strains were compared between the full consortium and the dropout community (N=9 each), p values are denoted as * < 0.05, ** <0.01, ***< 0.005.

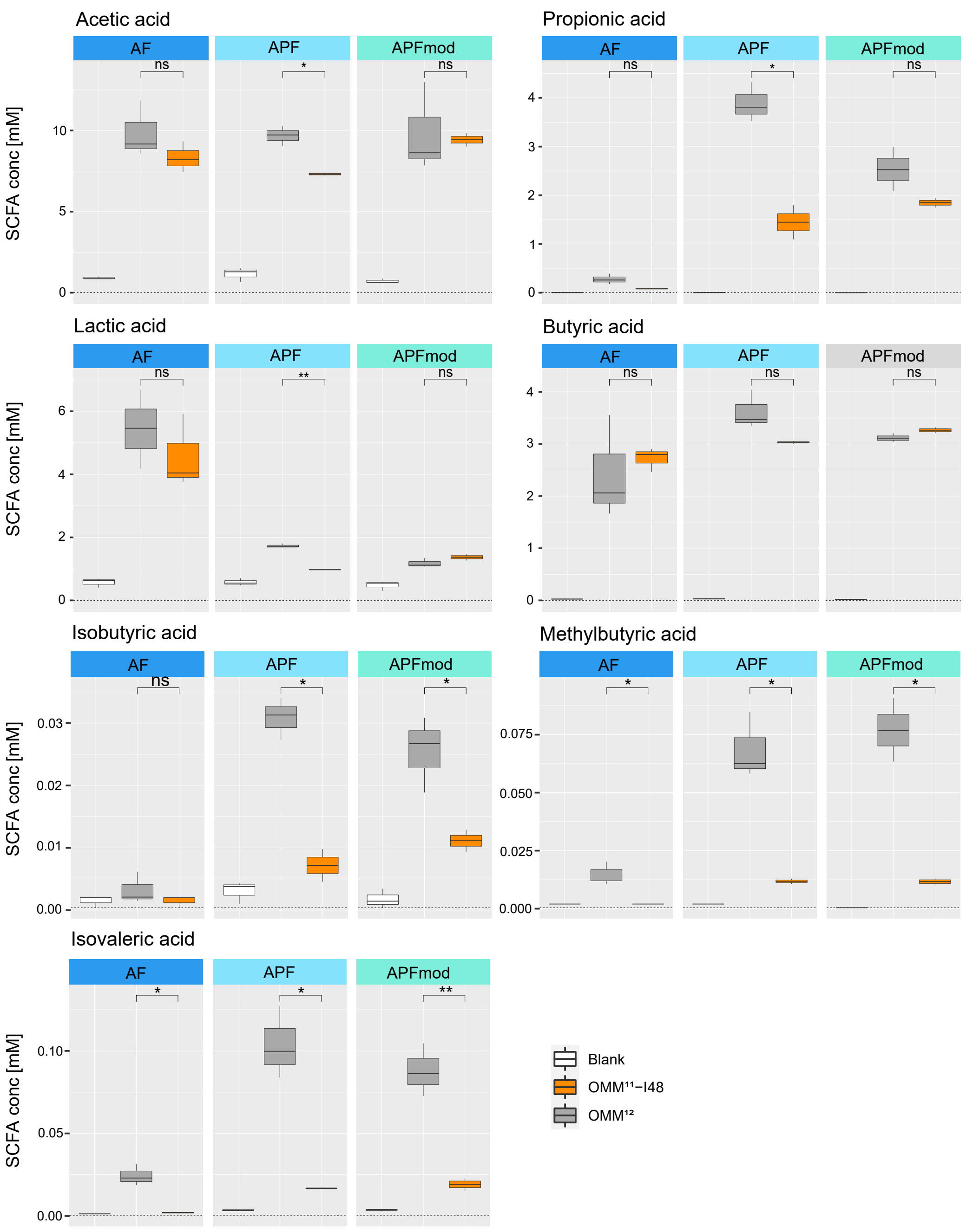


**Fig. S6. SCFA concentrations of *B. caecimuris* I48 dropout communities.** SCFA concentrations were determined by targeted metabolomics analysis of community spent media and fresh media (AF, APF and APF$^{mod}$ medium) and are shown as median with the corresponding upper and lower percentile. Using a t-test the SCFA concentrations in the *B. caecimuris* I48 dropout community were compared to the full consortium. p values are denoted as ns = not significant, * $<$ 0.05, ** $<$0.01.

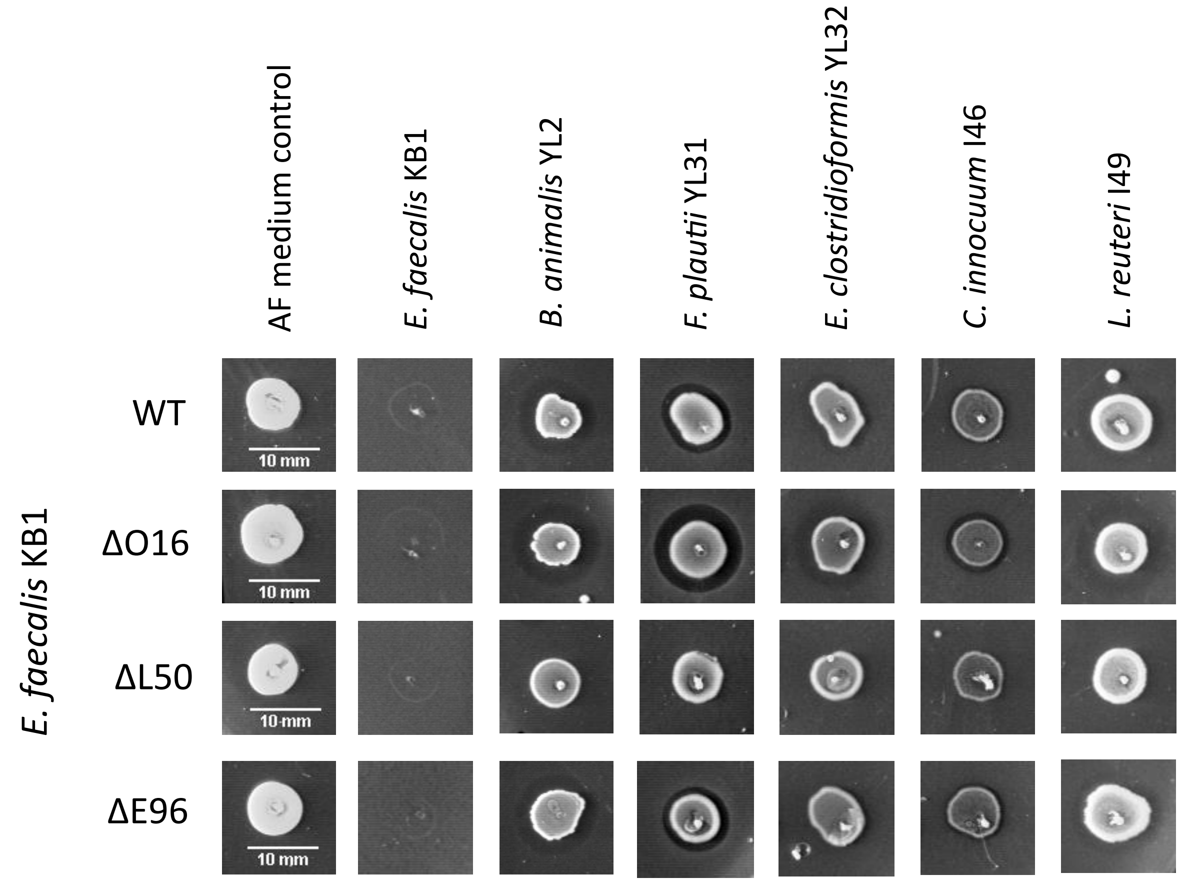


**Fig. S7. Phenotyping approach of *E. faecalis* KB1 wildtype and mutant strains.** Spot assays were used to test for the production of antibacterial compounds. *E. faecalis* KB1 wildtype and mutant strains lacking the loci for production of enterocin O16, enterocin L50 (A and B) and enterocin E96 were spotted onto a bacterial lawn of the initially susceptible strains (ref): *B. animalis* YL2, *F. plautii* YL31, *E. clostridioformis* YL32, *C. innocuum* I46 and *L. reuteri* I49; as well as on a lawn of *E. faecalis* KB1 wildtype and AF medium as control. The *E. faecalis* KB1 ΔL50 mutant strain was the only strain that did not show clear inhibition zones on any of the tested other strains, identifying enterocin L50 A and B as the inhibiting compound.

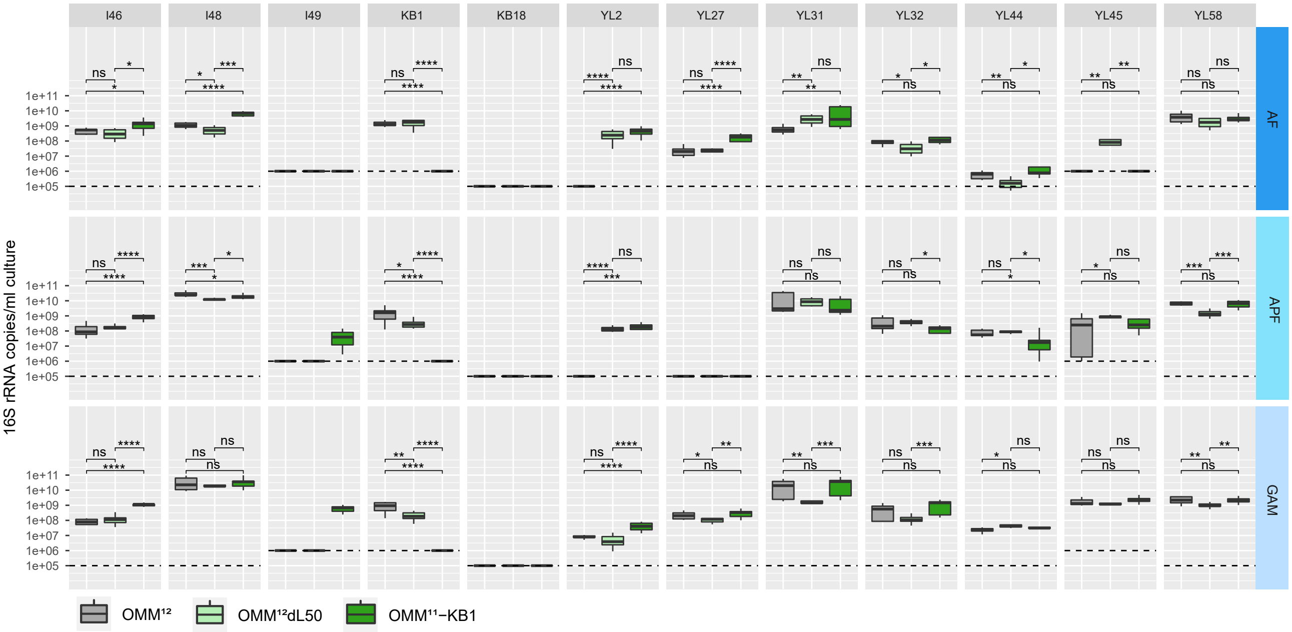


**Fig. S8. Influence of enterocin L50 on community assembly.** Absolute abundance of strains after four days of serial dilution in batch culture in AF, APF and GAM medium was determined by qPCR as normalized 16S rRNA copies per ml culture for the full consortium, a community including the *E. faecalis* KB1 ΔL50 mutant strain and a *E. faecalis* KB1 dropout community. Median absolute abundances are shown with the corresponding upper and lower percentile for all individual strains. Using a Wilcoxon test, absolute abundances of the individual strains were compared between the full consortium and the *E. faecalis* KB1 ΔL50 mutant strain community, or the dropout community (N=9 each), p values are denoted as ns = not significant, * $<$ 0.05, ** $<$0.01, ***$<$ 0.005.

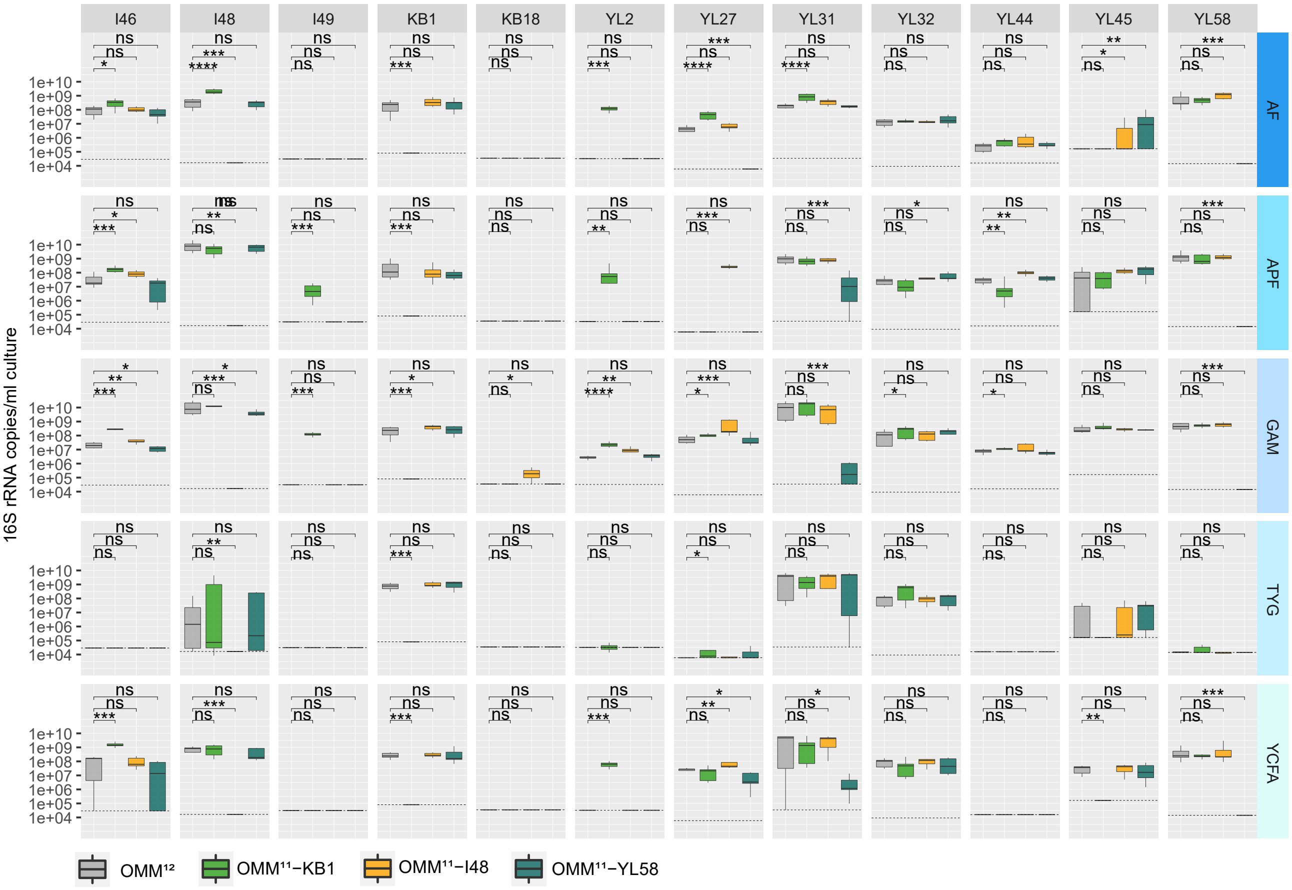


**Fig. S9. Community assembly of consortia lacking three keystone species in additional culture media.** Absolute abundance of strains after four days of serial dilution in batch culture in AF, APF, GAM, TYG and YCFA medium was determined by qPCR as normalized 16S rRNA copies per ml culture for the full consortium and communities lacking the three identified keystone species *E. faecalis* KB1, *B. caecimuris* I48, *B. coccoides* YL58. Median absolute abundances are shown with the corresponding upper and lower percentile for all individual strains. Using a Wilcoxon test, absolute abundances of the individual strains were compared between the full consortium and the corresponding dropout communities (N=9 each), p values are denoted as ns = not significant, * $< 0.05$, ** $<0.01$, ***$< 0.005$.

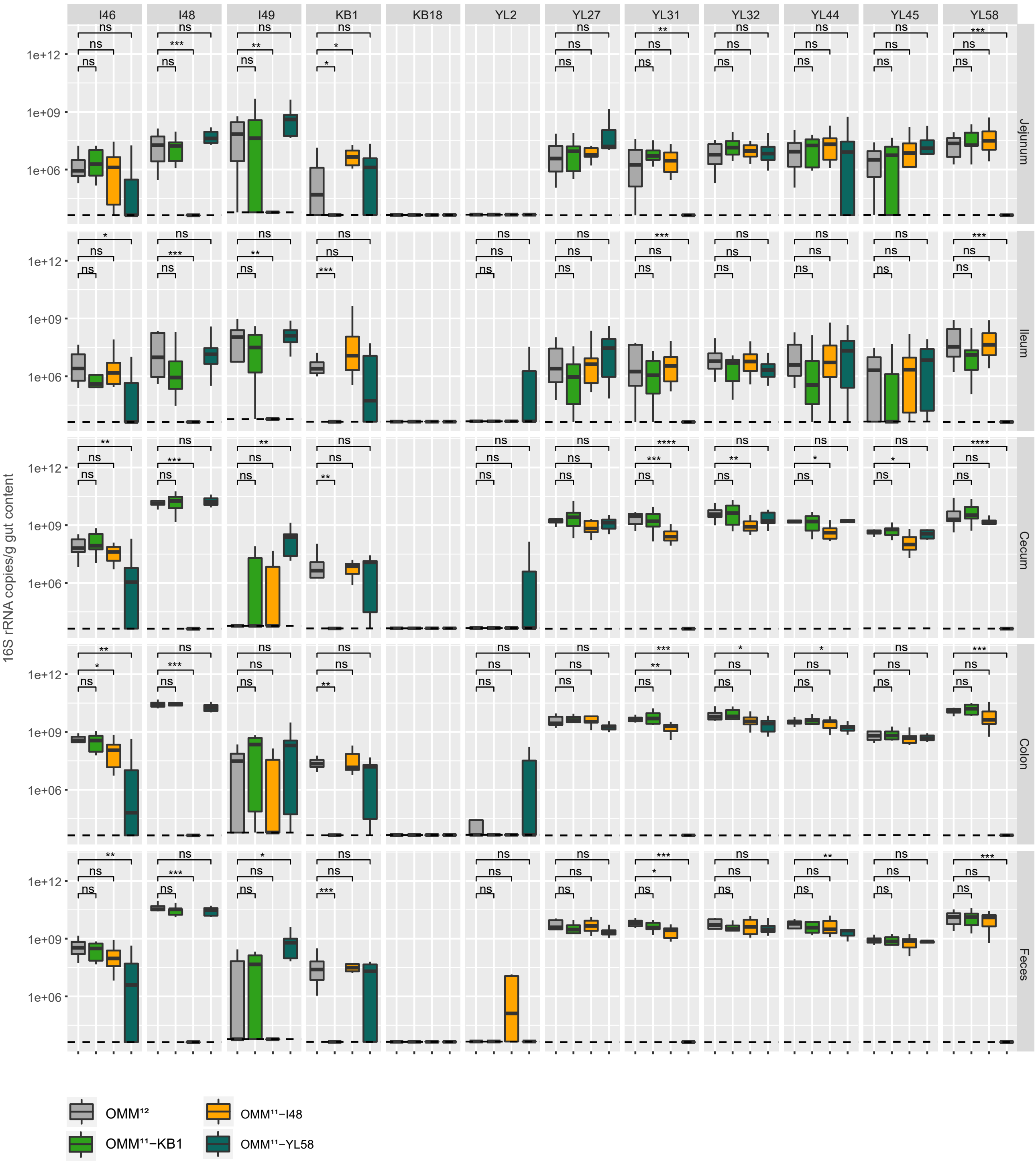


**Fig. S10. Community assembly across the different regions of the murine gut.** Absolute abundance of strains in the different regions of the murine gut (jejunum, ileum, cecum, colon and feces) was determined by qPCR as normalized 16S rRNA copies per g feces for the full consortium and communities lacking the three identified keystone species *E. faecalis* KB1, *B. caecimuris* I48, *B. coccoides* YL58. Median absolute abundances are shown with the corresponding upper and lower percentile for all individual strains (N=8-10 mice per group). Using a Wilcoxon test, absolute abundances of the individual strains were compared between the full consortium and the corresponding dropout communities, p values are denoted as ns = not significant, * < 0.05, ** <0.01, ***< 0.005.

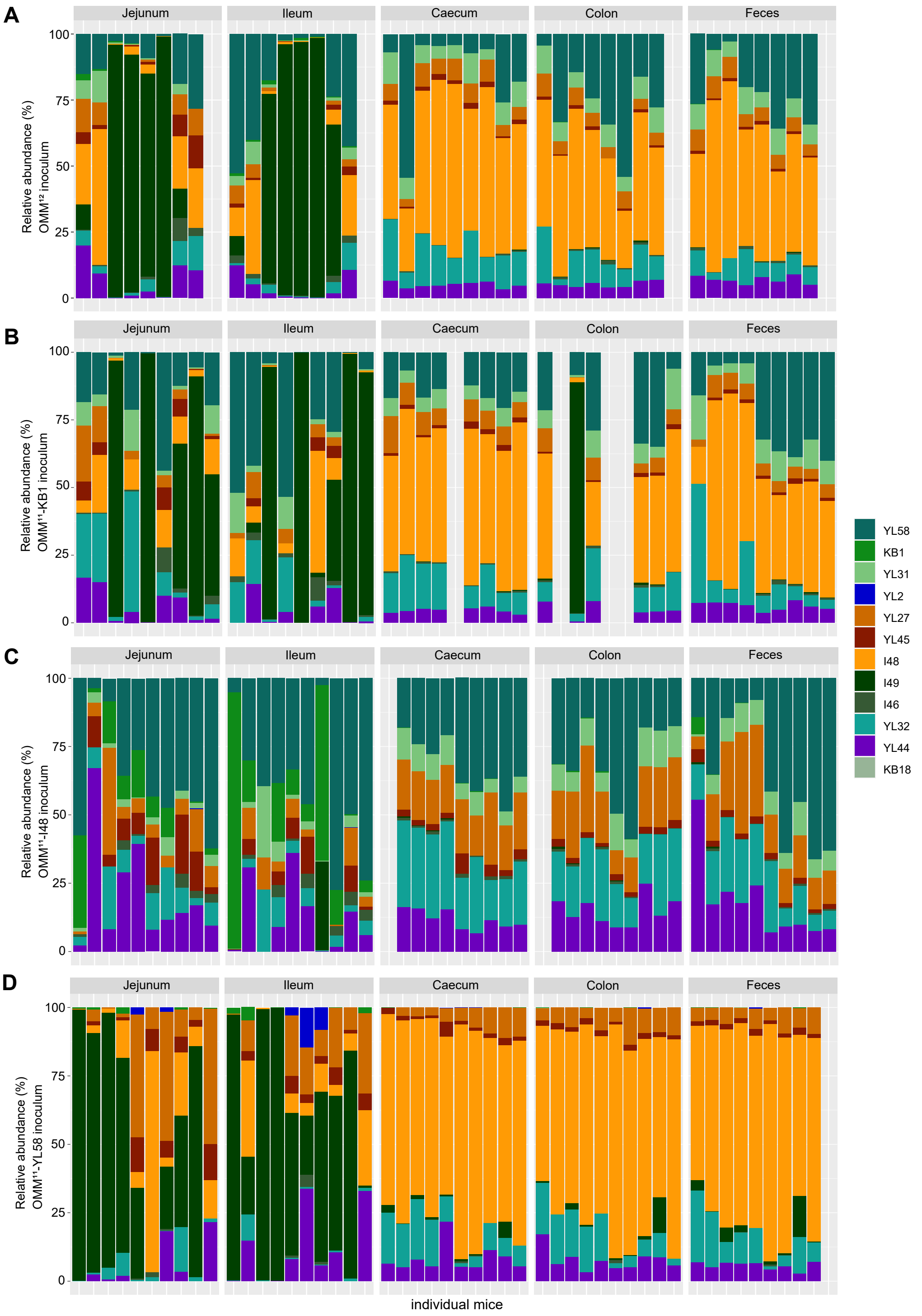


**Fig. S11. Relative abundances of communities across the different regions of the murine gut.** Based on absolute abundances of strains in the different regions of the murine gut (jejunum, ileum, cecum, colon and feces), the relative abundance profiles were determined for the full consortium and communities lacking the three identified keystone species *E. faecalis* KB1, *B. caecimuris* I48, *B. coccoides* YL58 for the individual mice (N=8-10 mice per group).

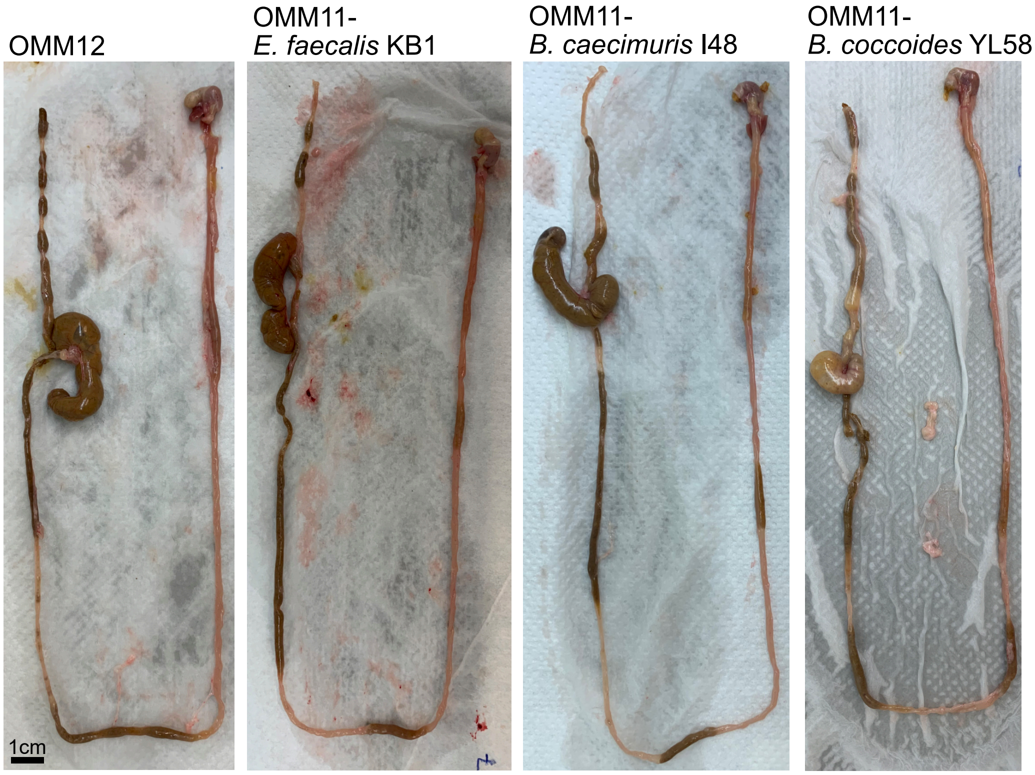


**Fig. S12. Physiology of GI tract of differently colonized mice.** Exemplary sections of the gastrointestinal tract of individual mice for each group, differing in their bacterial colonization (OMM$^{12}$, OMM$^{11}$-*E. faecalis* KB1, OMM$^{11}$- *B. caecimuris* I48, OMM$^{11}$-*B. coccoides* YL58). Strongest difference are visible in cecum size and texture.

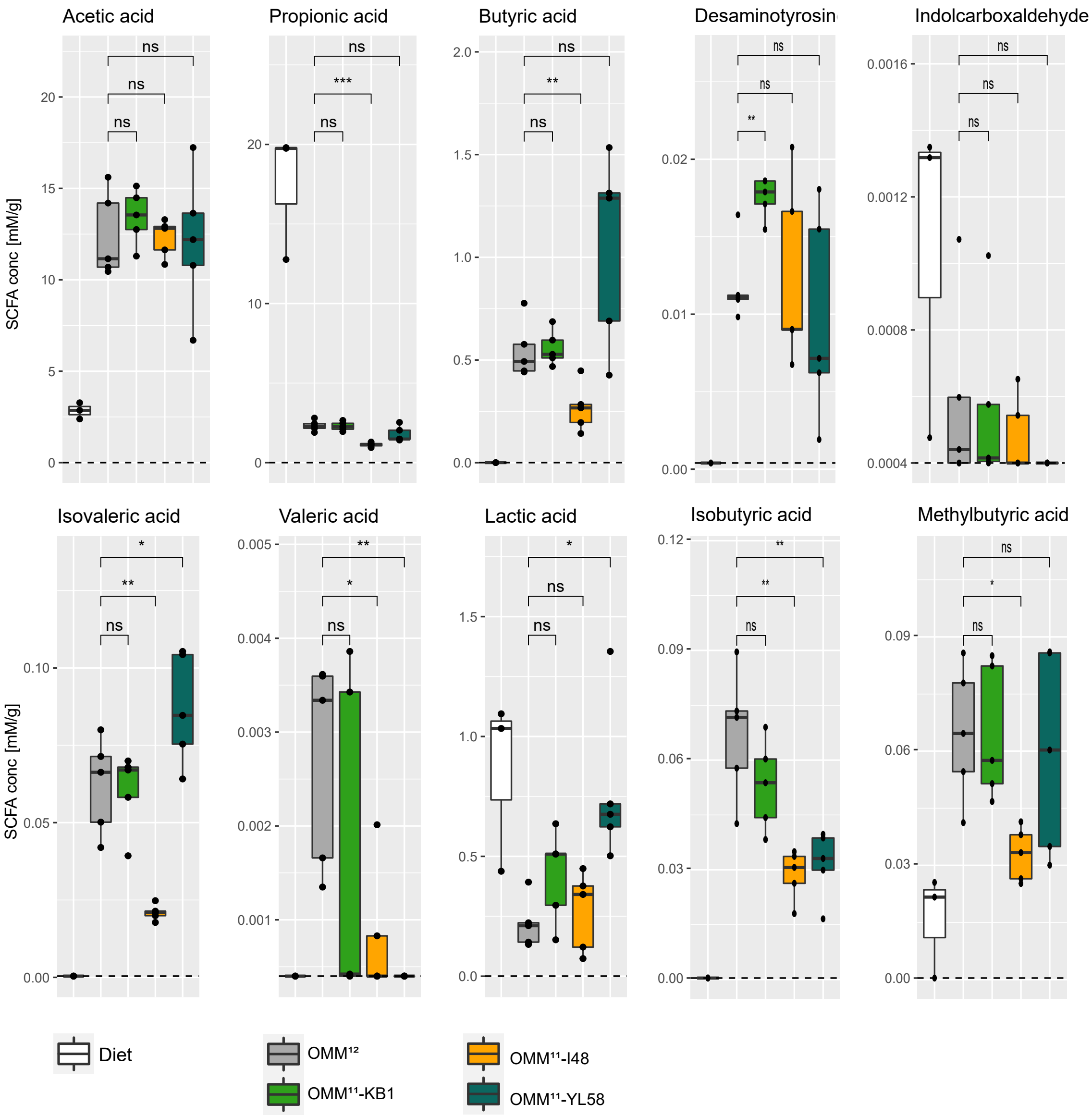


**Fig. S13. SCFA concentrations in the murine cecum.** SCFA concentrations were determined by targeted metabolomics analysis of cecal content of the differently colonized mice (N=5 per group) and are shown as median with the corresponding upper and lower percentile. Using a t-test the SCFA concentrations in the cecum of mice colonized with the full consortium were compared to the cecum content of mice colonized with the individual dropout communities (OMM$^{11}$-*E. faecalis* KB1, OMM$^{11}$- *B. caecimuris* I48, OMM$^{11}$-*B. coccoides* YL58), p values are denoted as ns = not significant, * < 0.05, ** <0.01.

#### Supplemental Tables

**Potential polysaccharide degradation enyzmes for inulin and xylan in *B. caecimuris* I48**

| | Locus Tag | Genome position | Annotation on NCBI | PUL database ID | Word search |
|---|---|---|---|---|---|
| Inulinase | I5Q79_16515* | 4035767-4037599 | DUF4980 domain-containing protein | GH32 in PUL22: 2,6-beta-D-fructofuranosidase | |
| Xylanase | I5Q79_08890* | 2096214-2098463 | endo-1,4-beta-xylanase | - | endo-1,4-beta-xylanase |
| | I5Q79_10285 | 2505152-2509216 | acetylxylan esterase | CE6 in PUL2: acetyl xylan esterase | - |
| | I5Q79_11165 | 2741693-2742679 | glycoside hydrolase family 43 protein | GH43_31 in PUL40: beta-xylosidase | - |
| | I5Q79_11420 | 2831509-2833095 | glycoside hydrolase family 30 protein | GH30_4/unk in PUL37: 1,4-beta-xylanase | - |
| | I5Q79_11750 | 2932024-2933343 | glycoside hydrolase family 43 protein | GH43_31 in PUL35: 1,4-beta-xylanase | - |
| | I5Q79_16930 | 4133515-4134651 | endo-1,4-beta-xylanase | - | endo-1,4-beta-xylanase |
| | * = Fig. Sx | | | | |

| Sources | Link | Last visit |
|---|---|---|
| *B. caecimuris* I48 genome | https://www.ncbi.nlm.nih.gov/nuccore/CP065319 | 4th October 2022 |
| PUL database cazy.org | http://www.cazy.org/PULDB/index.php?sp_name=Bacteroides+caecimuris+I48&sp_ncbi= | 4th October 2022 |

**Table S1. Screen for polysaccharide degradation enzymes in *B. caecimuris* I48.** The genome of *B. caecimuris* I48 was screened for polysaccharide utilization loci (PUL) specific for inulin and xylan degradation found in literature (ref) and a PUL database (Methods). Sequences of key enzymes for inulin and xylan degradation were blasted against the *B. caecimuris* I48 genome and names of key enzymes were checked in genome annotations of *B. caecimuris* I48 via word search ("1,4-beta-xylanase", "beta-xylosidase", "inulinase" and similar versions). Identified locus tags, annotations and PUL database IDs are listed.

| | AF | | APF | | GAM mod. (Himedia) | TYG | | YCFA | |
|---|---|---|---|---|---|---|---|---|---|
| component | amount per l | company | amount per l | company | amount per l | amount per l | company | amount per l | company |
| brain-heart infusion | 18,5 g | Oxoid | 18,5 g* | US Biological | - | - | - | - | - |
| trypticase soy broth | 15 g | Oxoid | 15 g* | US Biological | - | - | - | - | - |
| yeast extract | 5 g | Roth | 5 g | Roth | 2.5 g | 5 g | Roth | 2.5 g | Roth |
| peptone | - | - | - | - | 5 g | - | - | - | - |
| soya peptone | - | - | - | - | 3 g | - | - | - | - |
| proteose petone | - | - | - | - | 5 g | - | - | - | - |
| tryptone/pepton from casein | - | - | - | - | - | 10 g | Roth | - | - |
| casitone | - | - | - | - | - | - | - | 10 g | BD |
| meat extract | - | - | - | - | 2.2 g | - | - | - | - |
| liver extract | - | - | - | - | 1.2 g | - | - | - | - |
| digested serum | - | - | - | - | 10 g | - | - | - | - |
| heat-inactivated fetal calf serum | 3% (v/v) | Sigma-Aldrich | 3% (v/v) | Sigma-Aldrich | - | - | - | - | - |
| D-glucose | 0.5 g | Roth | - | | 0.5 g | 2 g | Roth | 2 g | Roth |
| 1:1 mixture of arabinose[a], fucose[b], lyxose[b], rhamnose[b], xylose[c] | - | - | 2.5 g | [a]Sigma-Aldrich, [b]TCI, [c]Roth | - | - | - | - | - |
| inulin | - | - | 2 g | Sigma-Aldrich | - | - | - | - | - |
| xylan | - | - | 2 g | Roth | - | - | - | - | - |
| starch | - | - | - | - | 5 g | - | - | 2 g | Roth |
| cellobiose | - | - | - | - | - | - | - | 2 g | Roth |
| mucin | - | - | 0.025% | Sigma-Aldrich | - | - | - | - | - |
| HCl-cysteine | 0.5 g | Sigma-Aldrich | 0.5 g | Sigma-Aldrich | 0.3 g | 0.5 g | Sigma-Aldrich | 1 g | Sigma-Aldrich |
| L-arginine | - | - | - | - | 1 g | - | - | - | - |
| L-trypthophan | - | - | - | - | 0.2 g | - | - | - | - |
| hemin | 1 mg | Sigma-Aldrich | 1 mg | Sigma-Aldrich | 0.01 g | - | - | 10 mg | Sigma-Aldrich |
| menadione | 0.5 mg | Sigma-Aldrich | 0.5 mg | Sigma-Aldrich | - | 1 mg | Sigma-Aldrich | - | - |
| vitamin K1 | - | - | - | - | 5 mg | - | - | - | - |
| hematin-histidine** | - | - | - | - | - | 1 ml | Sigma-Aldrich | - | - |
| trace element/vitamins*** | - | - | - | - | - | - | - | *** | *** |
| $K_2HPO_4$ | 2.5 g | Roth | 2.5 g | Roth | - | 100 ml 1M $K_xPO_4$ | Roth | 0.45 g | Roth |
| $KH_2PO_4$ | - | - | - | - | 2.5 g | - | - | 0.45 g | Roth |
| $MgSO_4$-$7H_2O$ | - | - | - | - | - | - | - | 0.09 g | Sigma-Aldrich |
| $Na_2CO_3$ | 0.4 g | Merck | 0.4 g | Merck | - | - | - | - | - |
| $NaHCO_3$ | - | - | - | - | - | | | 4 g | Sigma-Aldrich |
| NaCl | - | - | - | - | 3 g | - | - | 0.9 g | Roth |
| sodium thioglycollate | - | - | - | - | 0.3 g | - | - | - | - |
| TYG salt solution**** | - | - | - | - | - | 40 ml | - | - | - |
| $CaCl_2$ | - | - | - | - | - | 8 mg | Sigma-Aldrich | 0.09 g | Sigma-Aldrich |
| $FeSO_4$ | - | - | - | - | - | 0.4 mg | Sigma-Aldrich | - | - |
| resazurin | - | - | - | - | - | - | Sigma-Aldrich | 1 mg | Sigma-Aldrich |

| |
|---|
| * glucose-free |
| ** 12 mg hematin dissolved in 10 ml 0.2M histidine solution (pH 8) |
| *** biotin (0.01 mg/l, Sigma-Aldrich), cobalamin (0.01 mg/l, Sigma-Aldrich), folic acid (0.05 mg/l, Sigma-Aldrich), p-aminobenzoic acid (0.03 mg/, Sigma-Aldrich), pyridoxamine (0.15 mg/l, Fluka), thiamine (0.05 mg/l, Roth), riboflavin (0.05 mg/l, Sigma-Aldrich) |
| **** 0.05 g $MgSO_4$-$7H_2O$, 1 g $NaHCO_3$, 0.2 g NaCl in 100 ml $H_2O$ |

**Table S2. Overview of media compositions.** Comparison of composition of culture media used in this study. AF was used as described previously (Weiss et al, 2021). APF medium was first used in this study. Modified GAM medium is commercially available at Himedia (Himedia Labs). TYG and YCFA media were previously described (Whitacker et al., 2017 and Duncan et al., 2002)

PERMANOVA of Bray-Curtis dissimilarities
adjusted p-value (Benjamini-Hochberg)

| | OMM11-KB1 vs. OMM12 | OMM11-I48 vs. OMM12 | OMM11-YL58 vs. OMM12 |
|---|---|---|---|
| Jejunum | 0.6895 | 0.0099 | 0.4518 |
| Ileum | 0.4693 | 0.0648 | 0.4693 |
| Cecum | 0.512 | 0.0003 | 0.072 |
| Colon | 0.4502 | 0.00015 | 0.00015 |
| Feces | 0.9906 | 0.0003 | 0.0021 |

**Table S3. Statistical analysis of Bray-Curtis dissimilarities based on absolute strain abundances in the murine gut.** Bray-Curtis dissimilarities analysis of absolute strain abundances in the different regions of the murine gut was performed on samples obtained from mice colonized with the full consortium and communities lacking the three identified keystone species *E. faecalis* KB1, *B. caecimuris* I48, *B. coccoides* YL58 for the individual mice (N=8-10 mice per group). For each region, pairwise comparison of Bray-Curtis dissimilarities of the individual dropout consortia to the full consortium was performed using a permutational multivariate analyses of variance (PERMANOVA) in R using the function Adonis (method "bray" with 9,999 permutations). Obtained p values were adjusted using the Benjamini-Hochberg method. Significant values ($p < 0.05$) are highlighted in red.

| Primer name | Sequence | Amplification target |
| --- | --- | --- |
| P149 | gcaaattggtatttcacagtcc | L50A-L50B |
| P150 | catctaacaacctcttcttttattctt | |
| P151 | caaagacaacacgggataacactc | O16 |
| P152 | gtcgttaactttcacaaaatgtaagtc | |
| P153 | cgaatttttagtttcggctcttt | Ent96 |
| P154 | cgtttcaattaatgacctagacttc | |
| P159 | ggaaaacttcacgtatcggatc | Arm 1 L50A-L50B |
| P160 | ggtggtgtcgacaatatatctcctccaattattttttgttc | |
| P161 | ggtggtgtcgacttttatgatatattaaatttttaaagagactatga | Arm 2 L50A-L50B |
| P162 | ggtggtctgcaggtttgaaccagacctgcaat | |
| P163 | ggtggtggatccgatgattggtgggttagtagtagg | Arm 1 Ent96 |
| P164 | ggtggtgtcgacagtattatctcctttcgtcctctcc | |
| P165 | ggtggtgtcgacaaatttctaattagaataaccgtcctc | Arm 2 Ent96 |
| P166 | ggtggtctgcagtccgtaagggcgaattattta | |
| P167 | ggtggtggatccctcttggcttatcaaatttggaa | Arm 1 O16 |
| P168 | ggtggtgtcgacaaataaatccctactttctttttcctt | |
| P169 | ggtggtgtcgactgaatttaaaagatcatgttacgga | Arm 1 O16 |
| P170 | ggtggtctgcagcggtatatccttgctgagctttt | |
| pLT06_FW_b | cgtgtgctctacgaccaaaact | pLT06 vector |
| pLT06_RV_b | tcctcctttctattttgattag | |

**Table S4. List of primers for the construction of exchange vectors for the engineering of *E. faecalis* KB1.** Vector pLT06 was used for the deletion of enterocins L50A-L50B, Ent96 and O16 in *E. faecalis* strain KB1. DNA fragments of 500-1000 bp upstream and downstream of the gene targeted for deletion (homologous arms 1 and 2) were amplified by PCR, using primers to insert restriction sites. BamHI and SalI restriction sites were added to arms 1, and SalI and PstI sites were added to arms 2. The PCR products of the arms were digested using the appropriate restriction enzymes and ligated with pLT06.
